# A VTA GABAergic neural circuit mediates visually evoked innate defensive responses

**DOI:** 10.1101/493007

**Authors:** Zheng Zhou, Xuemei Liu, Shanping Chen, Zhijian Zhang, Yu-anming Liu, Quentin Montardy, Yongqiang Tang, Pengfei Wei, Nan Liu, Lei Li, Xiaobin He, Chen Chen, Guoqiang Bi, Guoping Feng, Fuqiang Xu, Liping Wang

## Abstract

Innate defensive responses are essential for animal survival and are conserved across species. The ventral tegmental area (VTA) plays important roles in learned appetitive and aversive behaviors, but whether it plays a role in mediating or modulating innate defensive responses is currently unknown. We report that GABAergic neurons in the mouse VTA (VTA^GABA+^) are preferentially activated compared to VTA dopaminergic (VTA^DA+^) neurons when a threatening visual stimulus evokes innate defensive behavior. Functional manipulation of these neurons showed that activation of VTA^GABA+^ neurons is indispensable for looming-evoked defensive flight behavior and photoactivation of these neurons is sufficient for looming-evoked defensive-like flight behavior, whereas no such role can be attributed for VTA^DA+^ neurons. Viral tracing and in vivo and in vitro electrophysiological recordings showed that VTA^GABA+^ neurons receive direct excitatory inputs from the superior colliculus (SC). Furthermore, we showed that glutamatergic SC-VTA projections synapse onto VTA^GABA+^ neurons that project to the central nucleus of the amygdala (CeA) and that the CeA is involved in mediating the defensive behavior. Our findings demonstrate that visual information about aerial threats access to the VTA^GABA+^ neurons mediating innate behavioral responses, suggesting a more general role for the VTA.

## INTRODUCTION

The ventral tegmental area (VTA) is a heterogeneous nucleus including dopaminergic (DA+) neurons, GABAergic (GABA+) and glutamatergic (Glut+) neurons (Dobi et al., 2010; Morales and Margolis, 2017; Yamaguchi et al., 2007), and its dysfunction has been implicated in depression (Nestler and Carlezon, 2006), schizophrenia (Davis et al., 1991), and addiction (Lüscher and Malenka, 2011). Dopamine neurons in VTA have been extensively studied for their role in reward and aversive processing (Bromberg-Martin et al., 2010; Fields et al., 2007; Schultz, 1998). It is widely accepted that VTA^GABA+^ neurons play an essential role in promoting aversion through inhibition of VTA^DA+^ neurons (Bocklisch et al., 2013; Jennings et al., 2013; Tan et al., 2012; van Zessen et al., 2012).

Recently, there has been an increase in evidence that demonstrates the role of GABA neurons in numerous behavioral and physiological processes, such as the involvement of GABA neurons in sensitizing dim-light vision in the retina (Herrmann et al., 2011), the acquisition of conditioned fear (Ciocchi et al., 2010; Haubensak et al., 2010) and predator odor evoked innate fear (Yang et al., 2016). In addition, it has been reported recently that dorsal raphe nucleus (DRN) GABA+ neurons are activated following looming stimulation (Huang et al., 2017) and Zona incerta (ZI) GABA+ neuronal projections to the periaqueductal gray (PAG) drive innate defensive responses (Chou et al., 2018) and also that VTA^GABA+^ neuronal projections to NAc enhance associative aversive learning (Brown et al., 2012).

In fact, the VTA also contributes to aversive cue processing, such as airpuffs, footshocks and free fall (Brischoux et al., 2009; Matsumoto and Hikosaka, 2009; Mirenowicz and Schultz, 1996; Wang and Tsien, 2011b). Consistent with these findings, VTA^GABA+^ neurons are evoked by footshock stimulation that induces conditioned place aversion (Tan et al., 2012). More importantly, it is of fundamental importance for animals across species to detect visual predatory-like environmental stimuli and generate avoidance behavior when necessary. Environmental stimuli require multiple sensory modality inputs that enable the animal to avoid potential threats (LeDoux, 2012). Given this, we speculate that the VTA is involved in processing visual, potentially life-threatening signals, and if so, what the underlying cell-specific neural circuitry mechanisms are.

VTA receives widespread inputs to encode multiple signals (Beier et al., 2015; Lammel et al., 2012; Morales and Margolis, 2017; Watabe-Uchida et al., 2012). The superior colliculus (SC), a retinal recipient structure, is a vital source for conveying visual signals to VTA neurons, implicated in detecting biologically salient stimuli (Dommett et al., 2005; Redgrave and Gurney, 2006). These findings hint that visual threatening signal, such as those deriving from predatory-like looming stimulus, may reach the VTA via SC, and that this pathway might play a role in processing of the innate defensive responses.

Selection and rapid execution of appropriate defensive responses, ranging from risk assessment, fighting, freezing to flight and attack, can be essential for survival when an animal faces imminent danger (Tovote et al., 2016; Ydenberg and Dill, 1986)A laboratory-based experimental paradigm has been established where an animal is exposed to an expanding dark disc (looming) stimulus to the upper visual field that mimics an approaching aerial predator (Yilmaz and Meister, 2013). This looming stimulus leads to innate defensive behaviors (e.g. flight-to-nest and hiding behaviors) (Yilmaz and Meister, 2013). This paradigm offers the opportunity to dig deeper into the neural circuitry underlying visually-evoked innate defensive behaviors (Evans et al., 2018; Huang et al., 2017; Li et al., 2018; Salay et al., 2018; Shang et al., 2018; Wei et al., 2015; Zelikowsky et al., 2018; Zhao et al., 2014).

Using this looming-evoked flight-to-nest behavioral paradigm with mice, we found that VTA^GABA+^ neurons were significantly activated by the aversive visual stimulus. Selective optogenetic inhibition and activation of VTA^GABA+^ neurons showed that they were indispensable and inducing for looming-evoked defensive behavior. Tracing and electrophysiological data show that VTA^GABA+^ neurons received glutamatergic inputs from SC and sent long projections to the CeA, which were also likely involved in the defensive behavior. To the best of our knowledge, this is the first evidence showing the involvement of VTA^GABA+^ neurons in an innate, evolutionally conserved, visually-evoked defensive responses.

## RESULTS

### 1. VTA^GABA+^ neurons respond to looming stimulus, which evokes defensive behavior

According to previous study (Yilmaz and Meister, 2013), mice were placed in an open field with a nest as a hiding place, and the presentation of an upper field expanding dark disc stimulus (looming stimulus) mimicking the approach of an aerial predator triggered transient intermittent periods of immobility (intersperse immobility) following by flight-to-nest and hiding in nest behavior (**Figure 1A**). A comparison of various looming stimuli, including front field expanding dark disc stimulus, upper field expanding white disc stimulus, lower field expanding dark disc stimulus and upper field expanding dark disc stimulus, only upper field expanding dark disc stimulus reliably triggered intersperse immobility following by robust flight-to-nest and hiding in nest behavior (upper field expanding dark disc stimulus evoked latency of onset of flight, 1.83 ± 0.21 sec; return to nest, 3.07 ± 0.33 sec; hiding time in 1 min after onset of looming stimulus, 71.86 ± 6.83 %; **Figure S3F**). In the following experiments, therefore, we used this type of looming stimulus, and henceforth refer to it as “looming”.

**Figure 1.**
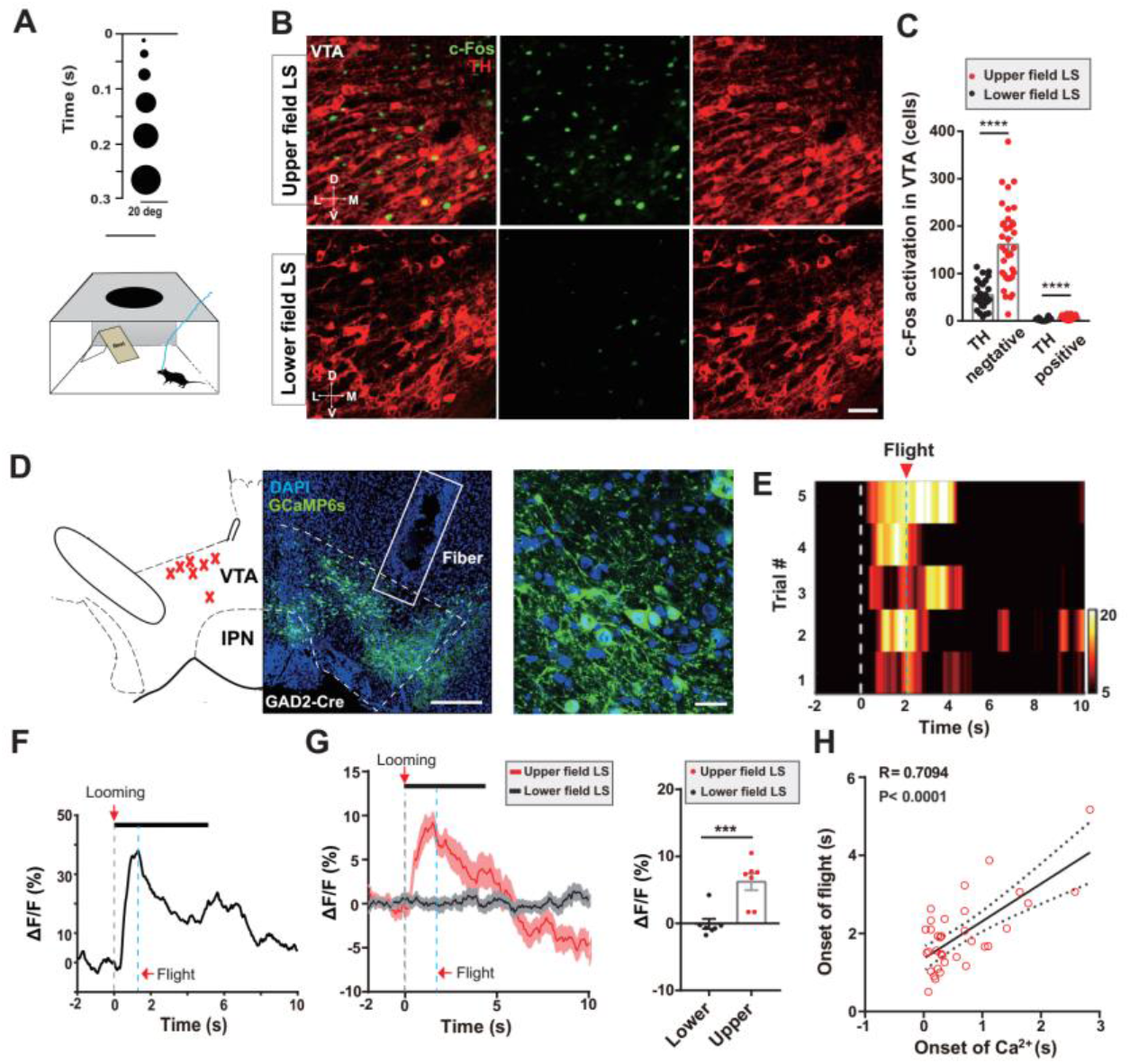
VTA^GABA+^ neurons respond to looming stimulus, which evokes defensive behavior. **(A)** Schematic paradigm of upper field looming stimulus (LS) in a nest-containing open field apparatus. **(B)** Representative images of c-Fos expression in the VTA following upper field LS (Top) and lower field LS (Bottom) control stimulus (green, c-Fos; red, TH; scale bars, 100 μm,). **(C)** Upper field LS led to higher c-Fos expression in VTA TH negative neurons compared to lower field LS (n=28-36 slices from 3 mice per group, for TH negative cells, *t*_62_=6.57, \*\*\*\**P*<0.0001; for TH positive cells, t62=4.909, \*\*\*\**P* <0.0001, Unpaired student test). **(D)** Left, schematic showing recording sites within the VTA; each red cross represents the optical fiber tip location from one mouse (n = 7 mice); Middle, representative image of AAV-*EF1α*::DIO-GCaMP6s expression in the VTA of GAD2-Cre mice (scale bars, 250 μm); Right, high-magnification image showing AAV-EF1a::DIO-GCaMP6s expression (scale bars, 20 μm). **(E)** Representative trial-by-trial heatmap presentation of calcium transients evoked by upper field LS in 5 trials from one mouse (white dotted line: onset of looming; blue dotted line: average latency of onset of flight). **(F)** Representative peri-event plot of the 1 trial of calcium transients (black bar represents presentation of the looming stimulus; gray dotted line: onset of looming; blue dotted line: latency of onset of flight). **(G)** Left, average calcium transients for the entire test group (red line, upper field LS; black line, lower field LS). Shaded areas around means indicate error bars, gray dotted line indicates onset of looming; Right, plot showing that there were more calcium transients during upper LS than lower field LS (n_Upper fleld LS_= 7 mice; n_Lower field LS_= 7 mice, *t_12_* =4.329, P=0.001; Unpaired student test). Gray dotted line: onset of looming; blue dotted line: average of latency of onset of flight from all mice. **(H)** Correlation analysis revealed that the onset of GCaMP6 transients were correlated with the onset of flight behavior ((n_Upper field LS_= 35 trials from 7 mice, linear regression *R*=0.7094, *F_1,33_* =33.44, \*\*\*\**P*< 0.0001). All data are presented as mean ± SEM.

We found that looming led to higher VTA c-Fos expression compared to that of a lower field looming stimulus control (**Figure 1B**). The distribution of c-Fos expression were located in the parabrachial and the paranigral portion of VTA. Tyrosine hydroxylase (TH) immunostaining revealed that the majority of c-Fos+ VTA neurons were TH-negative (**Figure 1C**, TH=3.90% vs Non-TH=96.1%). No significant increase of c-Fos expression was observed in rostromedial tegmental nucleus (RMTg) (**Figure S1**). Bear in mind that VTA is a heterogeneous nucleus and the largest neural population excluding dopaminergic neurons are GABAergic neurons (Dobi et al., 2010).

To confirm the recruitment of GABA+ neurons in VTA by looming stimulus and investigate the dynamics of such activation, we carried out *in vivo* calcium imaging in VTA by fiber photometry (Kim et al., 2016). Genetically encoded Ca^2+^ indicators (GCaMP6s) (**Figure 1D**) or GFP was expressed in VTA^GABA+^ neurons following stereotaxic infusions of the virus AAV-*EF1α*:: DIO-GCaMP6s into the VTA of the *GAD2*::Cre transgenic mice. VTA^GABA+^ neurons showed significant activation of GCaMP6s activity following upper field looming stimulus exposure (**Figure 1E, S2D and S2E**, 6.16%, ΔF/F mean), while no signal change in control mice expressing GFP (**Figures S2A–S2C**) or those exposed to control visual stimulus (**Figures S3B–S3E**). Calcium signal rose rapidly with the onset of looming (latency = 0.73±0.15 sec, mean ± SEM) and decayed slowly following looming stimulus offset (decay time constant = 3.19±0.20 sec) (**Figures 1G**). Calcium signal of VTA^GABA+^ neurons were active 1.1 sec precede onset of flight behavior (**Figures 1E and 1F**). Correlation analysis revealed that the onset of GCaMP6 transients signal were correlated with the onset of flight behavior (**Figures 1H**, linear regression R=0.7094, P < 0.0001). The temporal dynamics of the calcium signal correlated well with that of looming-evoked following by flight-to-nest behaviors.

These data demonstrate that VTA^GABA+^ neurons are robustly recruited by exposure to looming stimulus and suggest that they may be involved in mediating the defensive responses.

### 2. VTA^GABA+^ neurons mediate looming-evoked defensive behavior

To determine whether VTA has a role in looming-induced defensive behavior, we selectively suppressed neural activity in VTA^GABA+^ neurons using optogenetics. *Gad2::Cre* transgenic mice were bilaterally infected with AAV-*EF1α*:: DIO-eNpHR3.0-mCherry to express the light-activated chloride pump halorhodopsin (NpHR) selectively in VTA^GABA+^ neurons (**Figures 2A and 2B**). Delivery of consecutive yellow light to the VTA of these animals, but not those infected with control virus (AAV-*EF1α*:: DIO-mCherry), significantly suppressed defensive behavior elicited by looming stimulus, including an increased latency to return to nest, decreased speed of flight, and decrease in the percentage of hiding time spent in the nest after flight (latency: mCherry 4.6 ± 1.13 sec versus NpHR 27.7 ± 10.05 sec; speed: mCherry 1015 ± 310.7 % versus NpHR 220.5 ± 68.44 %; hiding time: mCherry 69.5 ± 7.54 % versus NpHR 25.17 ± 6.02 %; **Figures 2C and 2D**). These data suggest that neural activity in VTA^GABA+^ neurons is indispensable for defensive behavior to looming stimulus.

**Figure 2.**
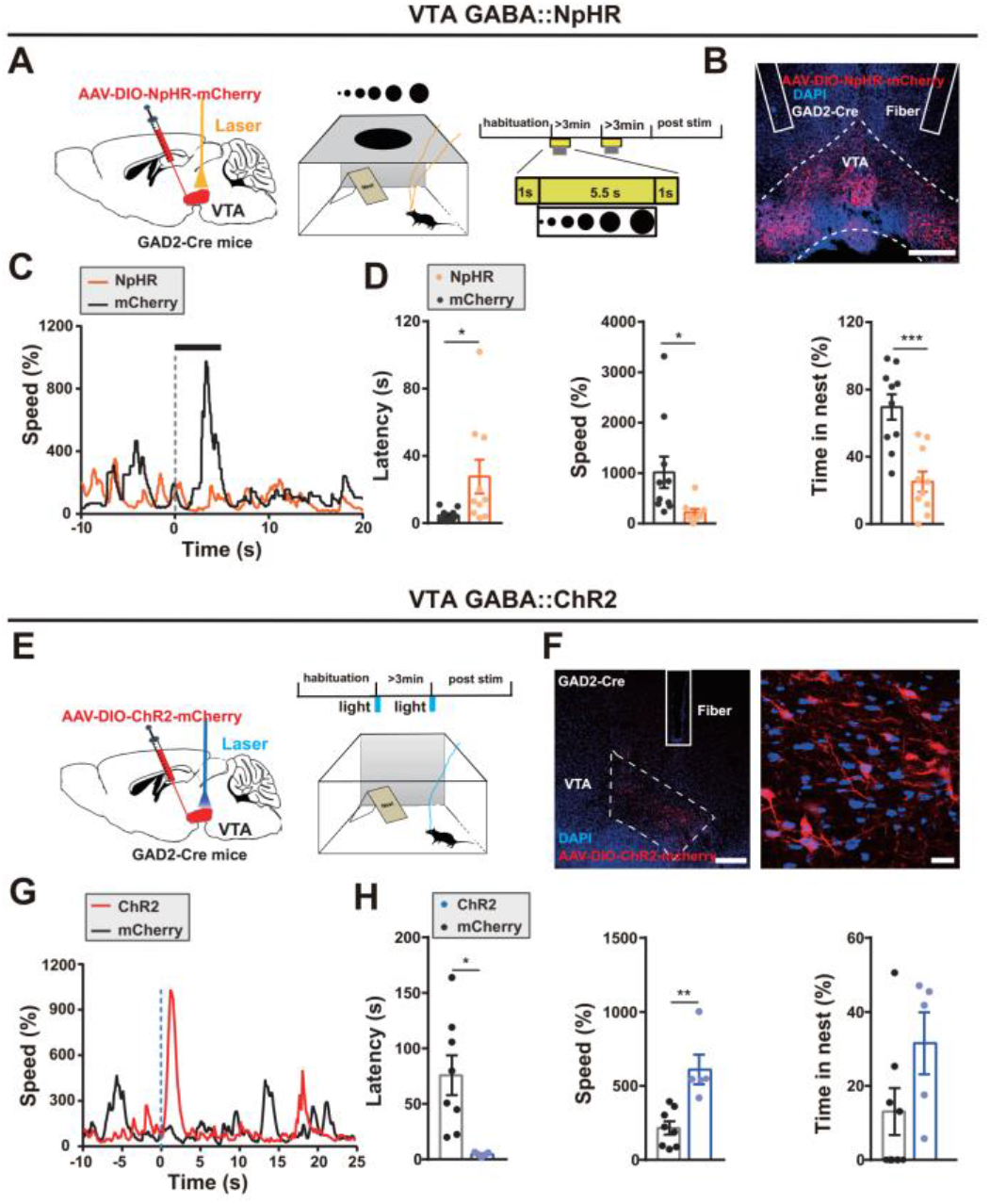
VTA^GABA+^ neurons mediate looming-evoked defensive behaviors. **(A)** Left, schematic diagram of bilateral optogenetic inhibition of VTA GAD2+ neurons during looming stimulus; Middle, open-field with nest looming apparatus; Right, looming test protocol. **(B)** Representative image showing NpHR virus expression in the VTA of a GAD2-Cre mouse and the position of the fiber track (blue, DAPI; red, NpHR-mCherry; scale bar, 250 μm; solid border lines, fiber tracks). **(C)** Representative curves show that bilateral inhibition VTA^GABA+^ neurons significantly decrease the instant speed compared with the control (blue dotted line: onset of looming stimulation). **(D)** Photoinhibition of VTA GAD2+ neurons resulted in higher latency back into the nest, lower speed, and shorter total percentage of hiding time spent in the nest after looming stimulus than mCherry controls (n_mCherry_ = 10 mice, n_NpHR_ = 9-10 mice, for latency, *t_18_*= 2.284, \**P*=0.0347; for speed, *t_17_*=2.372, \**P*=0.0297; for time in nest, *t_18_*=4.593, \*\*\**P*=0.0002; Unpaired student test). **(E)** Schematic diagram showing unilateral optogenetic activation of VTA GAD2+ neurons. **(F)** Representative image showing ChR2 virus expression in the VTA of a GAD2-Cre mouse and the position of the fiber track (blue, DAPI; red, ChR2-mCherry; scale bars, 20 μm, 250 μm respectively; solid border line, fiber track). **(G)** Representative curves show significantly instant speed evoked by opto-activation VTA^GABA+^ neurons compared with the control (blue dotted line: onset of blue light optical stimulation). **(H)** 2.5 s photoactivation of VTA GAD2+ neurons induced flight to nest behavior, shown by a decrease latency of flight-to-nest, an increase flight speed, and no change in total percentage of hiding time in the nest than mCherry controls (n_mCherry 2.5s_= 8 mice, n ChR2 2.5s=5 mice, for latency, *t_11_*= 3.085, \**P*=0.0104; for speed, *t_11_*=4.089, \*\**P*=0.0018; for time in nest, *t_11_*= 1.78, *P*=0.1028; Unpaired student test). All data are presented as mean ± SEM.

We then tested whether activation of VTA^GABA+^ neurons might trigger flight-to-nest behavior. *Gad2*::Cre transgenic mice were unilaterally infected with AAV-*EF1α*:: DIO-ChR2-mCherry to express the light-activated cation channel channelrodopsin (ChR2) selectively in VTA^GABA+^ neurons (**Figures 2E and 2F**). Mice were placed into the looming stimulus apparatus without the presentation of looming stimulus. Selective illumination of the VTA with blue light (5ms pulse, 60 Hz for 2.5 sec) elicited 1.29 ± 0.46 sec intersperse immobility following by flight-to-nest behavior in ChR2-expressing, but not control virus infected animals, and showed significantly decreased latency to return to nest, increased flight speed (latency: mCherry 75.75 ± 17.96 sec versus ChR24.46 ±0.82 sec; speed: mCherry 215.7 ± 44.9% versus ChR2 610.7 ± 100.7 %, **Figures 2G and 2H**). Longer blue light stimulation (5ms pulse, 60 Hz for 20 sec) induced flight-to-nest behavior and hiding time (mCherry 10.92 ± 6.98 % versus ChR2 58.43 ± 7.06 %, **Figures S4A and S4B**) ChR2 group. These results demonstrate that activation of VTA^GABA+^ neurons can induce innate defensive behaviors.

### 3. SC inputs to VTA mediate looming-evoked innate defensive behavior

To investigate possible upstream sources of visual inputs to VTA that might mediate looming stimulus responses, we examined mono-synaptic inputs to VTA^GABA+^ neurons using rabies virus tracing (Watabe-Uchida et al., 2012; Wickersham et al., 2007). *Gad2*::Cre mice were coinfected with AAV-*EF1α*:: DIO-RVG and AAV-EF1a:: DIO-TVA-GFP in the VTA and three weeks later infected with pseudo-typed and glycoprotein-deficient rabies virus (RV-EvnA-dG-dsRed) into the same coordinates (**Figures 3A and 3B**). After an additional week, mice were killed and examined for the distribution of upstreams of VTA^GABA+^ neurons. As expected, significant dsRed+ cells were identified in lateral habenula (LHb), PAG, and SC, regions known to provide inputs to VTA(Beier et al., 2015; Watabe-Uchida et al., 2012). For SC, dsRed+ cells were enriched mainly in the intermediate (IL) and deep layers (DL) (**Figures 3C and 3E**; mean RV+ / starter cell (%); SL, 0 %; IL, 63.65 %; DL, 56.95 %). Notably, dsRed+ cells were rare in the lateral geniculate nucleus (LGN) and primary visual cortex (V1) (**Figures 3D and 3G**, mean RV+ / starter cell (%); SC, 87.06 %; LGN, 0.061 %; V1, 0.236%) suggesting that SC provides the predominant source of direct inputs to VTA. Immunostaining revealed that the majority of VTA-projecting SC neurons in IL and DL were CaMKIIα+ (**Figure 3F**; mean ± SEM, 84.3%±3.23), suggesting that VTA^GABA+^ neurons receive direct monosynaptic CaMKIIα-positive inputs from the IL and DL layers of SC.

**Figure 3.**
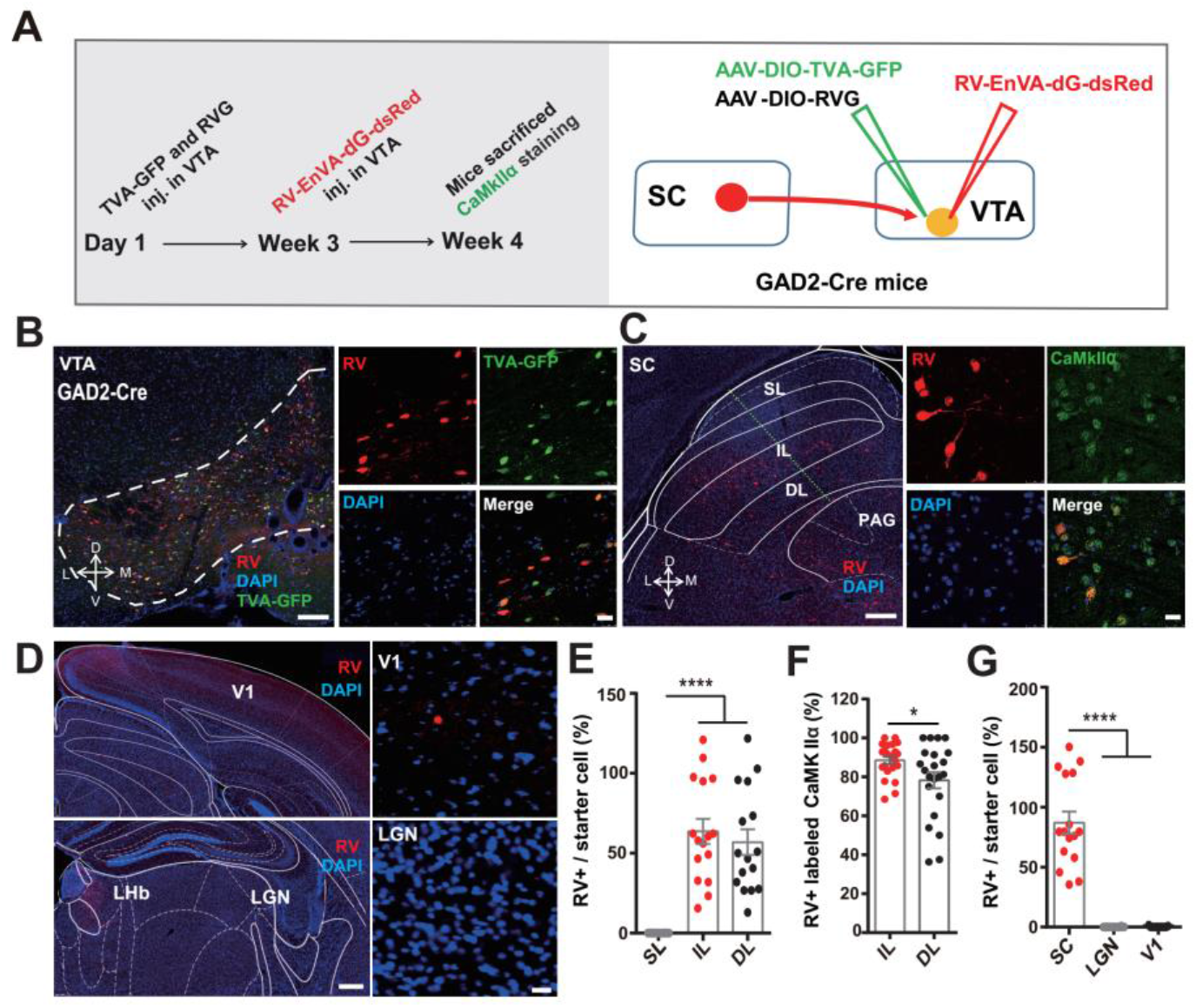
VTA^GABA+^ neurons receive direct monosynaptic CaMKIIα-positive inputs from the SC. **(A)** Schematic of rabies virus-based cell-type-specific monosynaptic tracing protocol. **(B)** Representative images denoting the starter cells in the VTA of GAD2-Cre mice (Red, rabies-dsRed; green, TVA; blue, DAPI, scale bar, 250 μm and 25 μm respectively). **(C)** Representative images showing retrograde labeling in the SC with inputs to VTA GAD2+ neurons, colabeled with CaMKIIα (Red, rabies-dsRed; green, CaMKIIα; blue, DAPI, scale bar, 250 μm and 25 μm respectively). **(D)** Representative images showing little or no rabies-dsRed signal from VTA GAD2+ neurons in other visual-related brain regions, V1 and LGN. **(E)** Plot showing rabies-dsRed positive neurons in SC shows that the intermediate and deep SC layers sent direct inputs to VTA^GABA+^ neurons (SL, Superior layer, IL, Intermediate layer, DL, Deep layer; scale bars, 100 μm and 25 μm respectively, n=16 slices from 4 mice, *F_2, 45_*= 28.51, \*\*\*\**P*<0.0001, one-way ANOVA). **(F)** IL and DL of SC neurons sending inputs to VTA^GABA+^ neurons, 84.3% ±3.23 were CaMKIIα-positive (n=22 slices from 4 mice, *t_42_* =2.254, \**P*=0.0294; Unpaired student test). **(G)** Plot showing rabies-dsRed positive neurons. SC sent many more direct inputs to VTA^GABA+^ neurons, rare signal was observed in other visual related brain regions, LGN and V1 (using starter cell number for normalization, n=11-16 slices from 4 mice, *F_2,38_*= 67.13, \*\*\*\**P*<0.0001, one-way ANOVA). V1, primary visual cortex; LGN, lateral geniculate nucleus; PAG, periaqueductal grey; LHb, lateral habenula. All data are presented as mean ± SEM.

Next, we investigated whether direct SC inputs to VTA might be functionally involved in looming-evoked defensive behavior. We selectively activated SC-VTA projections by delivering blue light (5ms pulse, 20 Hz for 2.5 sec) to the VTA in mice unilaterally infected in SC with *AAV-CaMKIIα*:: ChR2-mCherry (**Figures 4A and 4B**) and placed into the looming stimulus apparatus without the presentation of the looming stimulus. Photoactivation of this SC-VTA pathway induced 1.75 ± 0.36 sec intersperse immobility following by robust flight-to-nest and hiding in nest behavior, demonstrated by a significant increase in flight speed, a decreased latency to return to the nest, and an increased percentage of hiding time spent in the nest (latency: mCherry 31.53 ± 5.71 sec versus ChR2 4.12 ± 0.94 sec; speed: mCherry 212.5 ± 32.93% versus ChR2 451.5 ± 31.48 %; hiding time: mCherry 29.76 ± 8.32 % versus ChR2 81.02 ± 5.48 %; **Figures 4C and 4D** and **Video S1**) compared to the mCherry control group. Importantly, the defensive behavior was impaired by local pretreatment with the glutamate receptor antagonist (AP5/NBQX) into the VTA (**Figures S5A**) confirming a dependence of the behavioral effect on the activation of postsynaptic excitatory receptors in VTA. SC-VTA projections activation also elicited significant increases in heart rate and circulating corticosterone levels (**Figures S5B and S5C**), suggesting a widespread recruitment of downstream sympathetic arousal pathways. Direct photoactivation of CaMKIIα+ neurons in IL and DL of SC also resulted flight-to-nest behavior (**Figures S6A-S6E, S11A** and **Video S2**). These data suggest that activation of SC-VTA pathway can induce defensive responses in mice.

**Figure 4.**
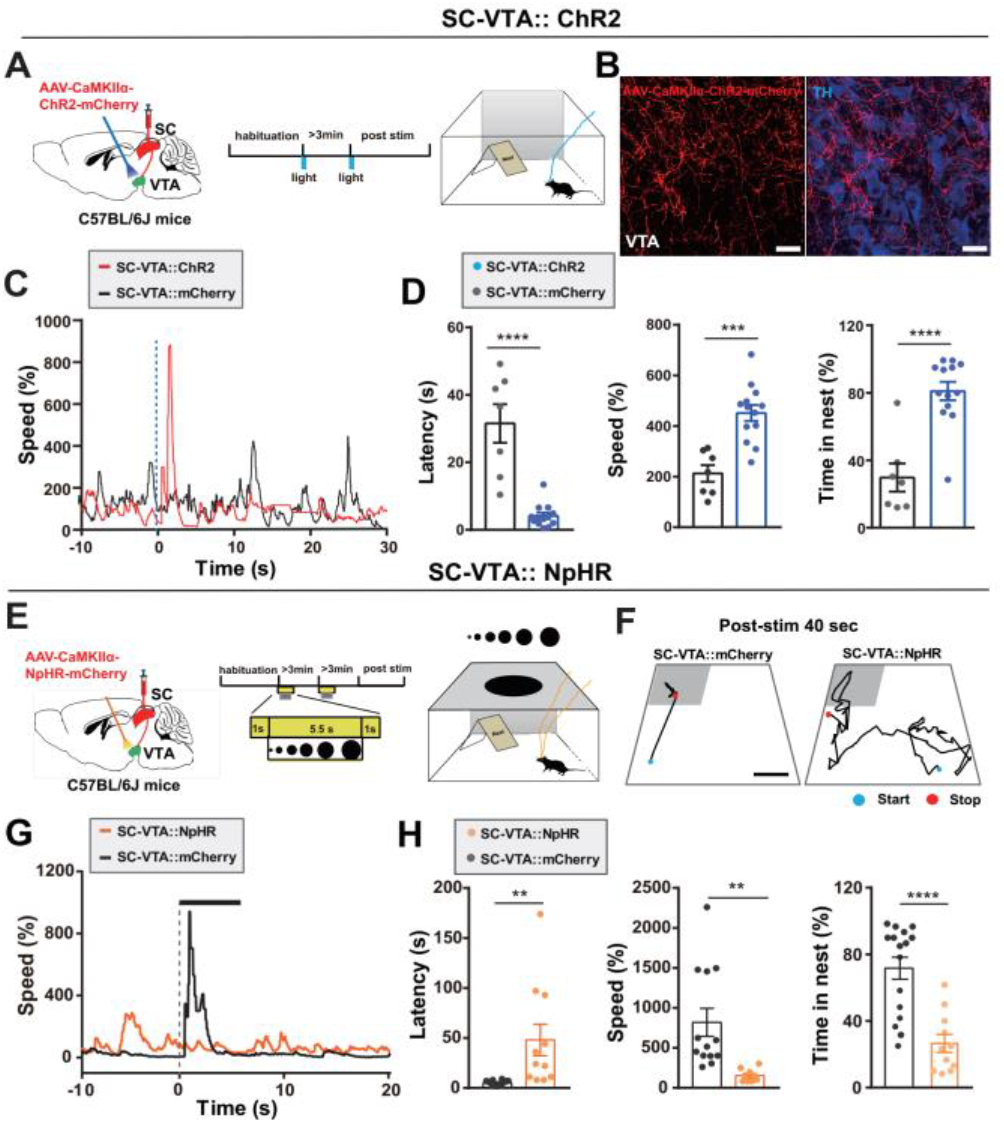
CaMKIIα:: SC-VTA pathway mediates looming-evoked defensive behavior. **(A)** Left, schematic showing unilateral blue light stimulation of CaMKII_αSC-VTA_:: ChR2; Middle, stimulation protocol; Right, open-field with nest looming apparatus. **(B)** Representative image showing ChR2 virus expression in the fibers from SC CaMKIIα-positive neurons in the VTA (red, AAV-CaMKIIα-ChR2-mCherry; blue, TH; scale bar, 25 μm). **(C)** Example of instant speed highlights the evoked flight behavior by opto-activation of CaMKIIαSC-VTA:: ChR2 compared with the mCherry control (blue dotted line: onset of blue light optical stimulation). **(D)** Photoactivation of CaMKIIα^SC-VTA^:: ChR2 resulted in flight to nest behavior, specifically, an increase in speed, shorter latency back into the nest, and higher total percentage of time spent in the nest after looming stimulus (n_mCherry_ = 7 mice, n_ChR2_=13 mice, for latency, *t_18_*= 6.394, \*\*\*\**P<0.0001;* for speed, *t_18_* =0.4834, \*\*\**P*=0.0001; for time in nest, *t_18_*=5.326, \*\*\*\**P<0.0001;* Unpaired student test). **(E)** Left, Schematic of bilateral optogenetic stimulation of CaMKIIα^SC-VTA^::NpHR, Middle, looming test protocol; Right, open-field with nest looming apparatus. **(F)** Representative motions tracks of two mice shows that optogentic activation of CaMKIIα_SC-VTA_:: NpHR resulted in less looming-evoked flight to the nest behavior in the open field with nest apparatus than that of mCherry controls. **(G)** Representative curves show the instant speed significantly decreased by light inhibition CaMKIIα^SC-VTA^ pathway compared with the mCherry control (gray dotted line: onset of looming stimulation). **(H)** Optogenetic activation of CaMKIIα^SC-VTA^::NpHR led to higher latency back into the nest, lower speed, and lower total percentage of time spent in the nest after looming stimulus (n_mCherry_= 13-16 mice, n_NpHR_= 9-11 mice, for latency, *t_25_*= 3.303, \*\**P*=0.0029; for speed, *t_25_* =3.121, \*\**P*=0.0054; for time in nest, *t_25_* =4.927, \*\*\*\**P<0.0001*; Unpaired student test). All data are presented as mean ± SEM.

We then examined whether the SC-VTA pathways is indispensable for looming stimulus evoked defensive responses. Mice were bilaterally infected with AAV-*CaMK2a*:: eNpHR 3.0-mCherry in SC (**Figures 4E** and **S11C**). Selective delivery of consecutive yellow light to the VTA of NpHR-expressing terminals, but not control virus infected animals, elicited a significant increase in latency to return to the nest, decreased the flight speed, and decrease in the percentage of time spent in the nest after looming stimulus exposure (latency: mCherry 4.91 ± 0.63 sec versus NpHR 48 ± 15.86 sec; speed: mCherry 818.8 ± 175.1 % versus NpHR 153.5 ± 26.82 %; hiding time: mCherry 71.67 ± 6.62 % versus NpHR 26.52 ± 5.37 %; **Figures 4F-4H** and **Video S3**). These data suggest that the CaMKIIα:: SC-VTA pathway is indispensable for the induction of innate defensive behavior by looming stimulus.

### 4. SC glutamatergic inputs activate VTA^GABA+^ neurons

To understand the impact of SC-VTA pathway activation by looming stimulus we performed *in vivo* and *in vitro* electrophysiology to examine neural activity in VTA-^GABA+^ neurons while activating SC-VTA inputs. *In vivo* multichannel extracellular recordings were carried out in mice following unilateral infection of SC with AAV-*CaMKIIα*:: ChR2-mCherry and selective delivery of blue light (5 ms pulse, 20 Hz) to the VTA. Among 97 VTA neurons recorded, spike width, firing rate, and burst firing characteristics allowed us to classify 35 as putative non-dopaminergic (non-DA+) (**Figure 5D**) and 62 as putative dopaminergic (DA+) neurons (**Figure S7B**), though we need to keep in mind that it has been reported that among the putative VTA^DA+^ neurons identified based on the classical electrophysiological characteristics, a minority could be non-dopaminergic neuron (Margolis et al., 2006; Ungless and Grace, 2012). For putative non-DA+ neurons, photostimulation of SC–VTA fibers resulted in time-locked firing with a mean latency of 5.65±3.44 ms (**Figure 5B**). The majority of neurons (25/35, 71.4%) exhibited an increase in firing rate, while only a small fraction (2/35, 5.7%) exhibited a decrease (**Figures 5C and 5D**). The short latency of the responses confirmed our retrograde tracing data that a majority of VTA^GABA+^ neurons receive direct excitatory SC inputs.

**Figure 5.**
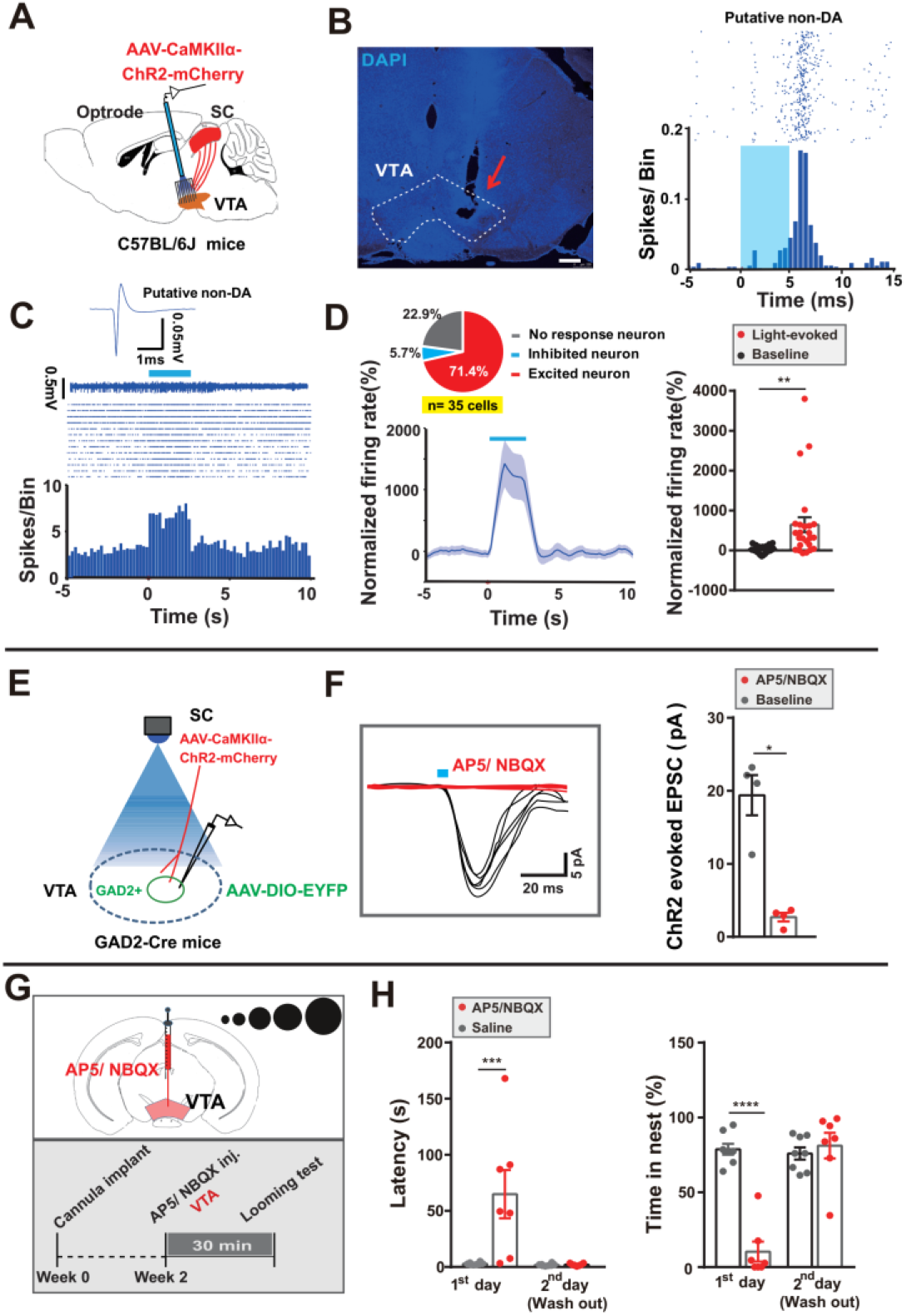
SC glutamatergic inputs activate VTA^GABA+^ neurons. **(A)** Schematic showing *in vivo* multichannel recording of single-unit VTA neuronal activity while optical stimulating CaMKIIα SC-VTA terminals. **(B)** Left, representative image showing the optrode position in the VTA (arrow, scale bar, 250 μm). Right, representative peristimulus time histogram (PSTH) and raster plot of a single neuron activated by CaMKIIα SC-VTA terminals stimulation, time-locked to 5 ms photostimulation (blue bar, terminals stimulation period). **(C)** Example PSTH and raster plot of a putative VTA non-DA neuron excited by CaMKIIα SC-VTA terminal optogenetic stimulation for a total period of 2.5 s. **(D)** Left, following terminal stimulation, 25/35 putative non-DA neurons (71.4%) were excited, 2 (5.7%) were inhibited, while 8 (22.9%) were unresponsive. Right, quantification of normalized firing rate of putative non-DA neurons (n= 35 units from 7 mice; *t_24_*=3.374, \*\**P*=0.0025, Paired student test). **(E)** Recording of evoked excitatory postsynaptic currents (eEPSCs) in VTA GAD2-positive neurons using patch-clamp slice recording during optogenetic stimulation of the CaMKIIα SC-VTA terminals. AAV-*EF1α*:: DIO-EYFP injections in GAD2-cre mice used to visualize GAD2-positive neurons. **(F)** EPSCs in VTA GAD2-positive neurons induced by CaMKIIα SC-VTA terminals light stimulation were not observed following injections of glutamate receptor antagonist (AP5+NBQX) (n= 4 cells from 3 mice, \**P*=0.0128, *t_3_*= 5.351, Paired t test). **(G)** Schematic of the looming behavioral paradigm after glutamate receptor antagonist (AP5+NBQX) or saline injections into the VTA. **(H)** Microinjection of glutamate receptor antagonist led to longer latency to return to nest and shorter percentage of time spent in the nest. There were no similar effects after drug washout. (n_saline_=8 mice, n_AP5+NBQX_=7 mice, Latency, day x drug effect interaction, \*\**P*=0.0091, *F_1,13_*= 9.355, bonferroni *post hoc* analysis, \*\*\**P*=0.0003, *P*>0.9999; Time in nest, day x drug effect interaction, \*\*\*\**P*< 0.0001, *F_1, 26_*=40.24, bonferroni *post hoc* analysis, \*\*\*\**P*<0.0001, *P*>0.9999, two-way ANOVA with bonferroni *post-hoc* analysis). All data are presented as mean ± SEM.

To better understand the functioning properties of SC inputs to VTA^GABA+^ neurons we carried out *in vitro* whole cell recordings in brain slices from *Gad2*:: Cre transgenic mice infected unilaterally with AAV-CaMKIIα::ChR2-mCherry in SC and AAV-*EF1α*:: DIO-EYFP in VTA (**Figure 5E**). Photostimulation of SC-VTA projections in the slice evoked excitatory postsynaptic currents (eEPSCs) in VTA^GABA+^ neurons that were eliminated by pretreatment with glutamate receptor antagonists (NBQX and AP5) (**Figure 5F**). To confirmed that glutamatergic signaling in VTA mediates looming-evoked responses, glutamate receptor antagonists (NBQX and AP5) were delivered via indwelling cannulas into the VTA 30 min before exposure to the looming stimulus (**Figure 5G**). Pre-treated animals showed a significant increase in the latency to return to nest and decrease in the percentage of time spent in the nest when compared to vehicle-treated controls or animals in which the drugs were allowed to washout (**Figure 5H**).

VTA^GABA+^ neurons play an important role in modulating the activity and responsivity of VTA^DA+^ neurons (Bocklisch et al., 2013; Jennings et al., 2013a; Nieh et al., 2016; Tan et al., 2012), primarily by directly inhibiting DA+ neuron firing. To examine whether such local inhibition might play a role in looming stimulus responses we carried out photostimulation of SC-VTA projections (20 Hz) while performing extracellular, single unit recordings of putative VTA^DA+^ neurons. Unexpectedly, we observed short latency (7.03±3.84 ms), time-locked firing of putative DA+ neurons following SC-VTA activation (**Figures S7A and S7C**). The majority of putative DA+ neurons (44/62, 71%) exhibited increased firing following stimulation (**Figures S7B, S7D and S7E**) and a minority (10/62, 16.1%) exhibited a decrease (**Figures S7B, S7F and S7G**).

Though the c-Fos activation of VTA^DA+^ neurons were rather weak under looming stimulus, considering the above electrophysiological recording data, it is intriguing to test the possible function of VTA^DA+^ neurons in looming-evoked defensive behavior. DAT::Cre mice were infected with AAV-*EF1α*:: DIO-ChR2-mCherry in the VTA (**Figures S7A-S7C**). Activation of VTA^DA+^ neurons did not elicit flight-to-nest behavior in open field without looming. We then examined whether the VTA^DA+^ neurons are indispensable for looming stimulus evoked defensive responses. DAT::Cre mice were bilaterally infected with AAV-*EF1α*:: DIO-eNpHR3.0-mCherry in the VTA (**Figure S8D**) and looming stimulus responses were measured during photoinhibition. Inhibition of VTA^DAT+^ neurons did not induce significant change in latency to return to nest or flight speed and increased the percentage time spent in the nest when compared with the mCherry mice (latency: mCherry 3.27 ± 0.52 sec versus NpHR 2.98 ± 0.47 sec; speed: mCherry 788.2 ± 187.8 % versus NpHR 947.9 ± 108.8 %; hiding time: mCherry 66.26 ± 7.35 % versus NpHR 88.34 ± 4.82 %; **Figures S8E and S8F**). There data proved the functions of VTA^GABA+^ neurons, but not VTA^DA+^ neurons, in looming-evoked flight-to-nest behavior.

Whole cell recordings in brain slices from TH::Cre mice in which photostimulation of SC-VTA projections was carried out confirmed the presence of evoked excitatory and inhibitory postsynaptic currents (eEPSCs and eIPSCs) in VTA TH+ neurons (**Figures S7H-S7K**). Notably, among 21 cells only 4 cells (19%) showed eIPSCs, confirming a relatively low probability of inhibition of DA+ neurons following SC-VTA pathway stimulation (**Figure S7I**).

For this part, our structural and functional results derived from viral tracing and electrophysiological recordings, respectively, show that VTA^GABA+^ neurons received functional glutamatergic inputs from SC, and that activating this pathway was inducing for acitivity of VTA^GABA+^ neurons.

### 5. CeA is a downstream target of the SC-VTA GABAergic pathway and it is involved in defense

Next, we sought to identify downstream targets of VTA^GABA+^ neurons that might mediate looming-evoked defense. Following infection of *Gad2*::Cre mice in VTA with AAV-*EF1α*:: DIO-mCherry (**Figures 6A and S9A**) dense labelled terminals were found in several brain regions, including the CeA, lateral hypothalamus (LH), LHb, and PAG (**Figures 6B, 6C and S9B-S9E**). Because CeA plays a critical role in mediating both learned and innate defensive behaviors (Gross and Canteras, 2012; Isosaka et al., 2015; LeDoux and Daw, 2018; Zelikowsky et al., 2018) we examined these projections in further details using tracing the relationship between input and output (TRIO) (Beier et al., 2015; Schwarz et al., 2015) and output-specific monosynaptic viral tracing (Gielow and Zaborszky, 2017).

**Figure 6.**
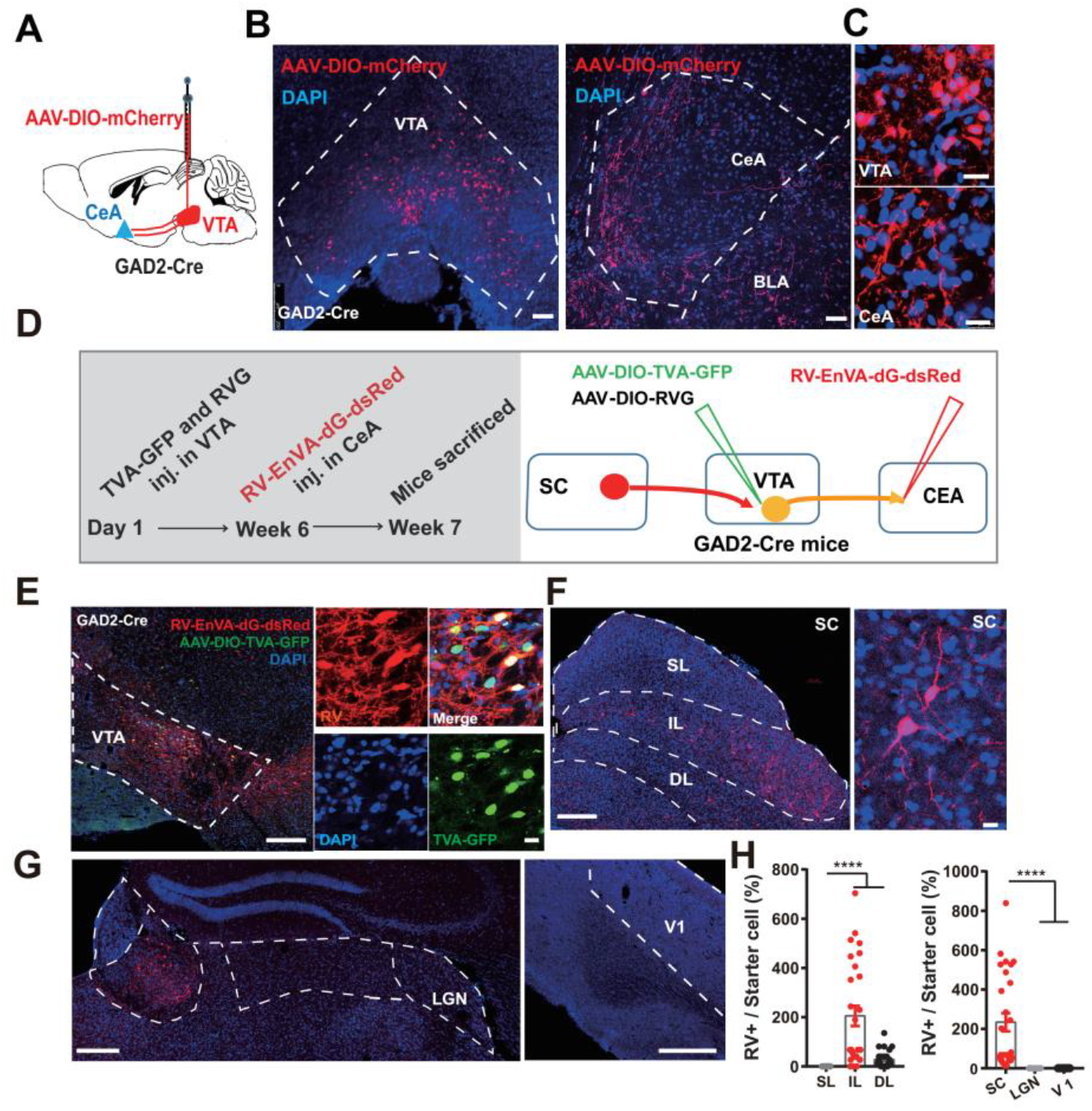
Output-specific monosynaptic viral tracing identifies the input-output relationship of VTA^GABA+^ neurons. **(A)** Schematic showing AAV-*EF1α*:: DIO-mCherry injection into the VTA of GAD2-Cre mice. **(B-C)** Representative coronal image showing targeted AAV-mCherry expression in the VTA and GAD2+ VTA-CeA terminals in the CeA (scale bar is 100 μm and is 10 μm respectively). **(D)** Schematic showing tracing of the input-output relationships between VTA^GABA+^ neurons: rabies-EnVA-dG-dsRed virus was injected into the CeA. AAV-*EF1α*:: DIO-RVG and AAV-*EF1α*:: DIO-TVA-GFP were co-injected into the VTA of the GAD2-cre mice. **(E)** Representative images showing VTA^GABA+^ starter cells (blue, DAPI, red, RV-EnVA-dG-dsRed, green, AAV-TVA-GFP, yellow, VTA GAD2+ starter cells, scale bars, 250 μm and 10 μm respectively). **(F-G)** Representative images showing substantial rabies-dsRed signal from CeA-projecting VTA^GABA+^ neurons in the SC (**F**: blue, DAPI; red, rabies-dsRed; IL and DL layers; scale bar, 250 μm and 10 μm respectively). No signal was observed in other visual related brain regions, LGN and V1 (**G:** blue, DAPI, red, RV-EnVA-dG-dsRed; scale bar, 250 μm). **(I)** Left, quantification of the number of rabies-dsRed labeled neurons in subregions of SC: the intermedial layer (IL), the deeper layer of the SC (DL) and the superficial layer (SL); (n =27 slices from 3 mice, data presented as mean ± SEM, \*\*\*\**P*<0.0001, *F_2,78_* = 18.5; one-way ANOVA). Right, quantification of the number of rabies-dsRed labeled neurons of the SC, V1, and LGN (n =14-27 slices from 3 mice, \*\*\*\**P*<0.0001, *F_2,60_* = 12.93; one-way ANOVA). All data are presented as mean ± SEM.

For TRIO, animals were infected unilaterally with CAV-Cre in CeA and AAV-*EF1α*:: DIO-histone-TVA-GFP, AAV-*EF1α*:: DIO-RVG, and EnvA-RV-dG-dsRed three weeks later in VTA (**Figure S10A**). Histological analysis revealed a dense distribution of dsRed+ cell bodies in the IL and DL layers of SC, while sparse labelling in the LGN and V1 (**Figures S10C, S10D and S10F**; mean RV+ / starter cell (%); SC, 400.8 %; LGN, 3.536 %; V1, 0 %). Tyrosine hydroxylase (TH) immunostaining showed that 78.4 ± 3.9 *%* of VTA starter cells were TH-negative (**Figures S10B and S10E**) suggesting the existence of a population of mainly non-DA+ VTA neurons that receive direct SC inputs and project to CeA. To confirm our findings we selectively examined the input-output connectivity of VTA^GABA+^ neurons using cell type-specific trans-synaptic rabies tracing (Gielow and Zaborszky, 2017). *Gad2*::Cre mice were infected with AAV-*EF1α*:: DIO-histone-TVA-GFP and AAV-*EF1α*:: DIO-RV-G) in VTA and EnvA-dG-RV-dsRed in CeA six weeks later (**Figures 6D and 6E**). Consistent with our TRIO results, histological analysis revealed frequent dsRed+ cell bodies in the IL and DL layers of SC and rare labeling in the LGN and V1 (**Figures 6E-6H**, mean RV+ / starter cell (%); SC, 234.1 %; LGN, 0 %; V1, 0 %).

Next, we attempted to confirm the existence of a functional VTA-CeA circuit using whole cell electrophysiological recordings in brain slices. *Gad2*::Cre mice were infected with AAV-*EF1α*:: DIO-ChR2-mCherry in VTA and possible blue light-evoked IPSCs responses were recorded in CeA (**Figures 7A and 7B**). In medial CeA (CeM), the majority of cells (7/10 cells, 70%) showed evoked IPSCs, while no evoked IPSCs were detected in lateral CeA (CeL, 0/21 cells, 0%) (**Figures 7C and 7E**). Importantly, evoked IPSCs were eliminated by pre-treatment of the slice with the GABA-A receptor antagonist bicuculline (**Figure 7D**) confirming that VTA^GABA+^ neuron inhibition of CeA is mediated by GABA.

**Figure 7.**
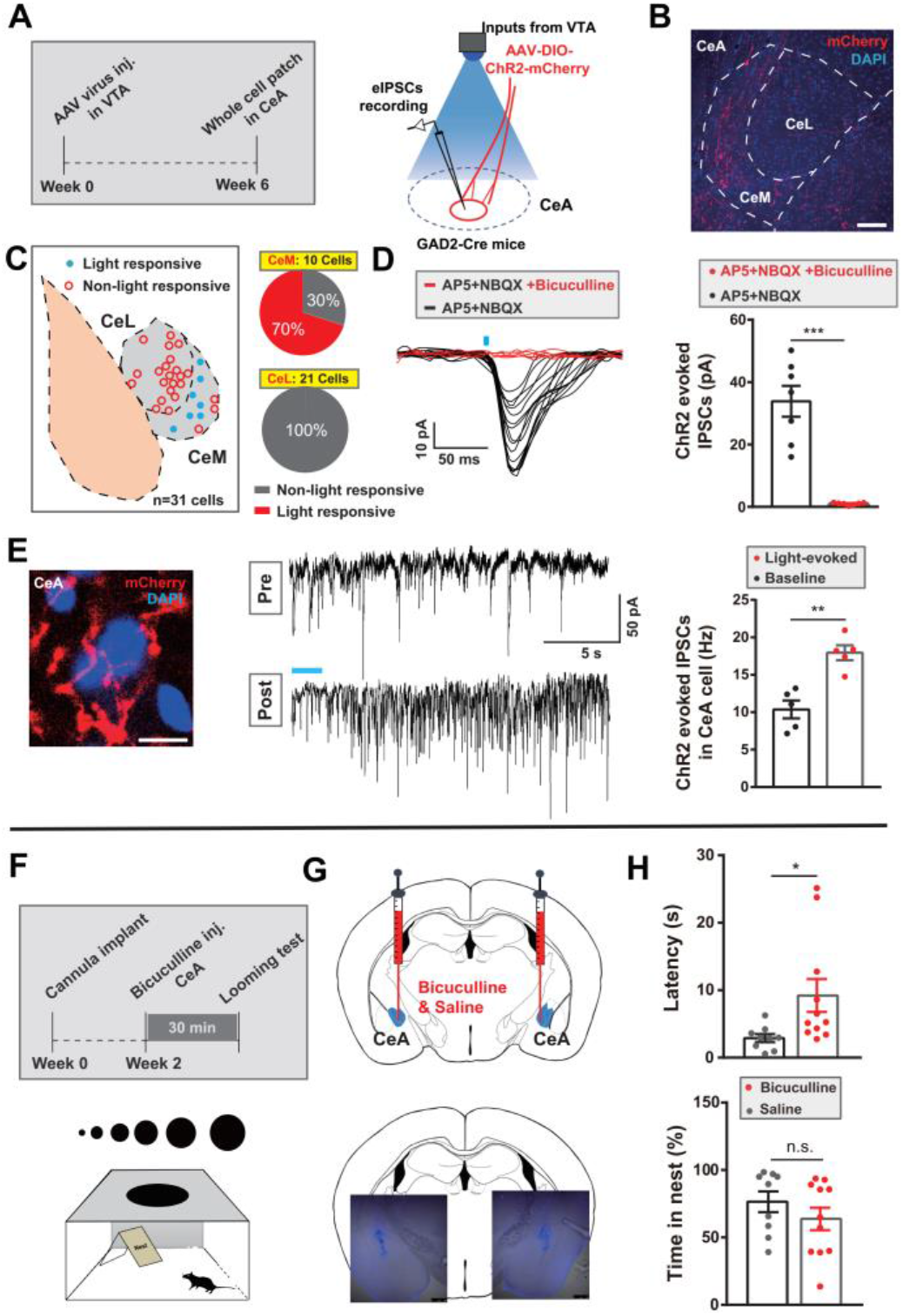
CeA as downstream target of SC-VTA GABA pathway is involved in looming-evoked flight-to-nest behavior. **(A)** Schematic showing timeline of recording of light-evoked inhibitory postsynaptic currents (eIPSCs) in CeA neurons using patch-clamp recording during optogenetic stimulation of GABAergic VTA-CeA terminals. **(B)** Representative coronal image showing the GAD2+ VTA-CeA terminals in the CeM subregion of CeA (scale bar is 100 μm). **(C)** Location of recordings in CeA. 21/31 cells recorded were located in the CeL and 10 cells located in the CeM. In CeM, eIPSCs were recorded in 70% cells (7 out of 10 cells) and 30% were non-responsive, while all the 21 cells in CeL were non-responsive. **(D)** GABA-A receptor antagonists, bicuculline abolished the IPSCs recorded in CeA induced by GABAergic VTA-CeA terminals optical stimulation (n= 7 cells from 6 mice, data presented as mean ± SEM, \*\*\**P*=0.0006, *t_6_*= 6.625, Paired student test) **(E)** Left, representative imagine of terminals in CeA; (red, terminals expressing with mCherry; blue, DAPI); Middle, light-induced increase of eIPSCs frequency from CeA neurons in brain slice patch clamp recording; Right, quantification of eIPSCs frequency (n=5 from 5 mice, \*\**P*=0.0055, *t_4_*= 5.451, Paired student test). **(F-G)** Top, schematic experimental timeline; Bottom, looming apparatus used following GABA-A receptor antagonist bicuculline or saline bilateral injections into the VTA. **(H)** Microinjection of GABA receptor antagonist bicuculline resulted in higher latency to return to nest, and no change in the percentage of time spent in the nest); (n_saline_= 9 mice, n_bicuculline_= 11 mice; for latency, *t_18_*=2.305, \**P*=0.0333; for time in nest, *t_18_*=1.101, *P*=0.2856; Unpaired student test). All data are presented as mean ± SEM.

To test whether inhibition of CeA plays a function role in looming-evoked defense we delivered the GABA-A receptor antagonist bicuculline directly into the CeA via indwelling cannulas 30 minutes before exposure to looming stimulus (**Figures 7F and 7G**). Bicuculline treated animals showed a significant increase in latency to return to nest and no significant change the hiding time in nest when compared to vehicle treated control animals (**Figure 7H**) arguing for a critical role of inhibition in CeA in visual-evoked defensive responses.

In summary, these data from tracing, slice recording and pharmacological experiments indicate that VTA^GABA+^ inhibitory inputs to CeA are high likely functionally involved in looming-evoked defense.

## DISCUSSION

In the current study, we found that exposure to a predator-like looming stimulus was associated with a rapid activation of VTA^GABA+^ neurons and that this is both indispensable and inducing for looming stimulus evoked defensive responses. Neural tracing experiments revealed that the circuit responsible for this behavior includes SC to VTA and then on to CeA, suggesting that VTA^GABA+^ neurons serve as a critical conduit in the processing of behavioral responses to aversive visual information.

The superior colliculus (SC) is sensitive to unexpected biologically stimuli, and has a functional role in detecting transient, rather than static, visual features (Redgrave and Gurney, 2006). A recent study has identified that the deep layers of the medial superior colliculus (dmSC) encode visual threat stimuli (Evans et al., 2018), which were normally recognized as unexpected sensory events. The pathway from SC-VTA that processes unexpected biological saliency (Dommett et al., 2005) has been identified using electrophysiological recording and rabies tracing (Dommett et al., 2005; Watabe-Uchida et al., 2012). Our viral tracing data together with functional studies, including *in vivo* multichannel recording and acute brain slice recording, complements this previous work and adds clarity by directly demonstrating that VTA-^GABA+^ neurons receive monosynaptic glutamatergic inputs from IDSC (Figure 3), which includes dmSC.

We identified that the SC-VTA pathway mediates innate defensive behaviors evoked by visual threat. We found that photostimulation of DLSC glutamatergic terminals of this population that project to VTA induces transient interspersed immobility following by flight-to-nest and hiding-in-nest behavior, which mimics the behavioral responses associated with looming-evoked defense.

Previous studies have shown that VTA GABA neurons receive inputs from subcortical structures, such as SC, LHb and PAG (Beier et al., 2015; Morales and Margolis, 2017). Aversive stimuli can induce activation of bed nucleus of the stria terminalis (BNST) projections to VTA, and these can drive activation of VTA GABA+ neurons (Jennings et al., 2013a). In our study, we found that VTA^GABA+^ neurons mediate looming-evoked innate defensive responses through the glutematergic pathway from IDSC to VTA. Fiber photometry recording revealed that VTA^GABA+^ neurons became active following looming stimulus before flight and also during flight. Inhibition of VTA^GABA+^ neurons significantly suppressed flight-to-nest behavior elicited by the looming stimulus, while activation of VTA^GABA+^ neurons also triggered transient interspersed immobility followed by flight-to-nest behavior, similar to the behavior following activation of the glutematergic pathway from SC to VTA. According to the economics hypothesis of flight from predators proposed by Ydenberge et al, prey may be aware of the predator well before it decides to flight. Premature flight may actually lead to vulnerability since it may increase salience and draw the attention of the predator (Ydenberg and Dill, 1986). The transient interspersed immobility seems like “risk assessment-like” behavior, which may have a suitable temporal window to allow for the detection, evaluation and preparation for making a decision to flight from the predator, all of which depended on the contextual factors, such as the existence of the escape routes (Blanchard et al., 2011; Evans et al., 2018; Tovote et al., 2016). Taken together, our data suggest that survival in a threatening environment activates the glutamatergic pathway from SC to VTA^GABA+^ neurons, which is involved in processing visual threats. VTA^GABA+^ neurons could potentially process the incoming visual signal inputs from SC and generate the adaptive behavioral responses to threats based on saliency and motivational value. One major contribution of this study is the finding that VTA^GABA+^ neurons are involved in processing potentially life-threatening innate fear signals.

Unexpected salient visual cues elicit an increase in the firing rate of DA+ neurons (Bromberg-Martin et al., 2010; Horvitz, 2000; Schultz, 1998) and visually evoked responses of VTA^DA+^ neurons depend on inputs from the intermediate and deep layer of SC (Dommett et al., 2005). In our c-Fos experiment, we observed that VTA^GABA+^ neurons were preferentially activated by looming stimulus, but a minority of VTA^DA+^ neurons were activated (Figure 1C). However, we did not observe that activation of VTA^DA+^ neurons induced interspersed immobility, flight-to-nest and hiding-in-nest behavior. Local GABA+ neurons are known to modulate DA+ neuron excitability and firing (Bocklisch et al., 2013; Jennings et al., 2013a; Nieh et al., 2016; Tan et al., 2012). Notably, 2.5 s photoactivation of VTA^GABA+^ neurons can induce flight-to-nest behavior but does not affect hiding time in the nest compared with looming-evoked behaviors. On the contrary, 20 s photoactivation of VTA^GABA+^ neurons that induced fight-to-nest behavior also led to longer hiding time in nest compared to that with 2.5 s photoactivation. This phenomenon may be due to increasing GABAergic excitation that results in stronger recruitment of inhibition of VTA DA, which may contribute to the modulation of hiding-in nest behavior. In evidence of this possibility, our data revealed that inhibition of VTA^DA+^ neurons during looming stimulation results in prolonged hiding time in nest. In line with this finding, recent work shows that inhibition of VTA DA neuronal activity during footshocks enhances fear (Luo et al., 2018).

Furthermore, data following administration of GABA receptor antagonists in the CeA during looming stimulus show that interspersed immobility behavior is still present, following by reduced latency of flight-to-nest, but no observable hiding time in nest. Our data suggest that the CeA, a downstream target of VTA^GABA+^ neurons, is involved in the flight-to-nest behavior evoked by looming. Most importantly, this data implies the involvement of distributions of VTA long-range projecting GABA neurons in flight behavior, and local VTA^GABA+^ neurons in hiding behavior via inhibition of VTA^DA+^ neurons. Indeed, our *in vivo* and *in vitro* electrophysiological recording data shows that minority of VTA^DA+^ neurons were inhibited following stimulating CaMKIIα SC-VTA terminals.

In our study, we posit that inhibition of local VTA^GABA+^ neurons on VTA^DA+^ neurons might be involved in regulation of looming induced defense responses. An interesting experiment would be to characterize the electrophysiology of the interaction of VTA-^GABA+^ neurons and VTA^DA+^ neurons during looming evoked defensive responses in freely moving animals.

VTA^GABA+^ neurons are heterogeneous subpopulations that include local GABA neurons and long-range GABA projection neurons (Morales and Margolis, 2017). Accumulating reports reveal roles of long-projecting GABAergic neurons in shaping behavioral outputs, such as feeding (Jennings et al., 2013b), avoidance (Lee et al., 2014), conditioned defensive behavior (Tovote et al., 2016) and innate defensive behavior (Chou et al., 2018). In addition, the role of VTA long-projecting GABAergic neurons in enhancing stimulus-outcome learning has been reported (Brown et al., 2012).

Next, we explored the downstream targets of VTA^GABA+^ neurons. Several brain regions, including the CeA, LH, LHb, and PAG had terminals that projected from VTA-^GABA+^ neurons. Given that the CeA plays a critical role in mediating both learned and innate defensive behaviors (Gross and Canteras, 2012; Isosaka et al., 2015; LeDoux and Daw, 2018; Zelikowsky et al., 2018), it was intriguing to test the possible function of VTA projecting GABA neurons to CeA.

We examined these projections in further details using tracing the relationship between input and output (TRIO) (Beier et al., 2015; Schwarz et al., 2015) and output-specific monosynaptic viral tracing(Gielow and Zaborszky, 2017). Our viral tracing data demonstrated that VTA^GABA+^ neurons sent long projections to CeA and these VTA^GABA+^ neurons received monosynaptic inputs from IDSC.

Furthermore, acute patch clamp slice recording shows that the VTA^GABA+^ projections preferentially form functional connectivity with CeM, the major output of the CeA that mediates defensive behaviors output (Ciocchi et al., 2010; Haubensak et al., 2010; LeDoux et al., 1988; Tovote et al., 2016), but not CeL. Recent reports show that CeA is involved in conditioned flight behavior, and that a competitive inhibitory circuit in CeA facilitates the selection of freezing and flight behavior where inhibition of CeM leads to flight (Fadok et al., 2017; Yu et al., 2016,Tovote, 2016). This flight behavior likely originates downstream of CeA in regions such as PAG and hypothalamus (Fadok et al., 2017; Tovote, 2016; Gross, 2012). It is possible that looming-activated VTA-^GABA+^ neurons send long inhibitory projections to the CeM subregion, which would inhibit CeM locally, thereby promoting flight behavior over freezing behavior. Our results are consistent with previous studies (Isosaka et al., 2015; Kalin et al., 2004; Salay et al., 2018), indicating the involvement of CeA in flight behavior evoked by innate threats, suggesting possible overlapping neural circuit bases for innate threats and learned threat processing in the amygdala (Gross and Canteras, 2012; LeDoux and Daw, 2018). Given the accumulating reports on amygdala involvement in defensive behaviors, including this current study, it would be interesting to further investigate the mechanism underlying the relation of innate threats and learned threat processing.

The selection and rapid execution of an appropriate defensive response, including the quick detection of the salient environmental cue, successfully flight to safety and hiding, require the integration of multiple sensory information inputs, spatial memory, and also requires adaptation to the environment. IDSC plays an important role in integrating multiple sensory inputs from environmental stimuli to rapidly escape from potential danger (Stein et al., 2009). Pharmacological infusion and electric stimulation of IDSC generate broad spectrum of defensive responses, including orienting, flight and freezing (Redgrave et al., 1981; Sahibzada et al., 1986). Very recently, Evans et al. identified that dmSC neurons that send projections to dPAG encode threat stimuli and when their firing reaches a synaptic threshold at dPAG, escape behavior is initiated (Evans et al., 2018). In our study, activation of IDSC glutamatergic neurons, which includes this part of dmSC, and IDSC-VTA pathway contributed to transient interspersed immobility, then flight-to-nest and hiding in nest behavior. The dmSC activity predicted the decision to flight 0.9 sec before onset of flight and dPAG neurons start to response only after onset of flight (Evans et al., 2018). Our results show that VTA^GABA+^ neurons were activated by looming stimulus before flight (latency of looming-evoked onset of Ca^2^+ signal of VTA^GABA+^ neurons: 0.73 sec; latency of looming-evoked onset of flight 1.83 sec) and sustained activity during flight.

The responses of VTA^GABA+^ neurons before flight initiation likely indicated the response to threat inputs through the pathway from IDSC to VTA before initiation of the appropriate behavior. This evidence fits with the time course of the transient interspersed immobility response to looming stimulus. Bearing in mind the potential gating role of GABA in filtering incoming inputs (Ren et al., 2012), our data suggests that the IDSC to VTA^GABA+^ pathway carries visual threat inputs. Understanding how animal access the risk and generate the most optimized option for defense through intrastimuli competition based on saliency and motivational value would be a very interesting future direction.

Our previous study demonstrated that glutamatergic neurons in mILSC projected to an LP-LA circuit mediating looming-evoked freezing behavior in an open-field without nest (Wei et al., 2015). Our current neural tracing results confirmed that CeA projecting VTA^GABA+^ neurons receive the inputs from the IDSC. This study provides a potential anatomical basis for further research to disentangle the behavioral selection processes that occur within the amygdala during unconditioned- and conditioned-defensive responses.

A detailed mechanistic understanding of the neural basis of these circuits will provide new insights to the potential mechanisms of survival across species, as well as the maladaptive behavior in fear- and anxiety-related mental disorders (Deisseroth, 2014; Lüthi and Lüscher, 2014; Pitman et al., 2012; Tovote et al., 2015).

## Supporting information

**Supplementary figure 1.**
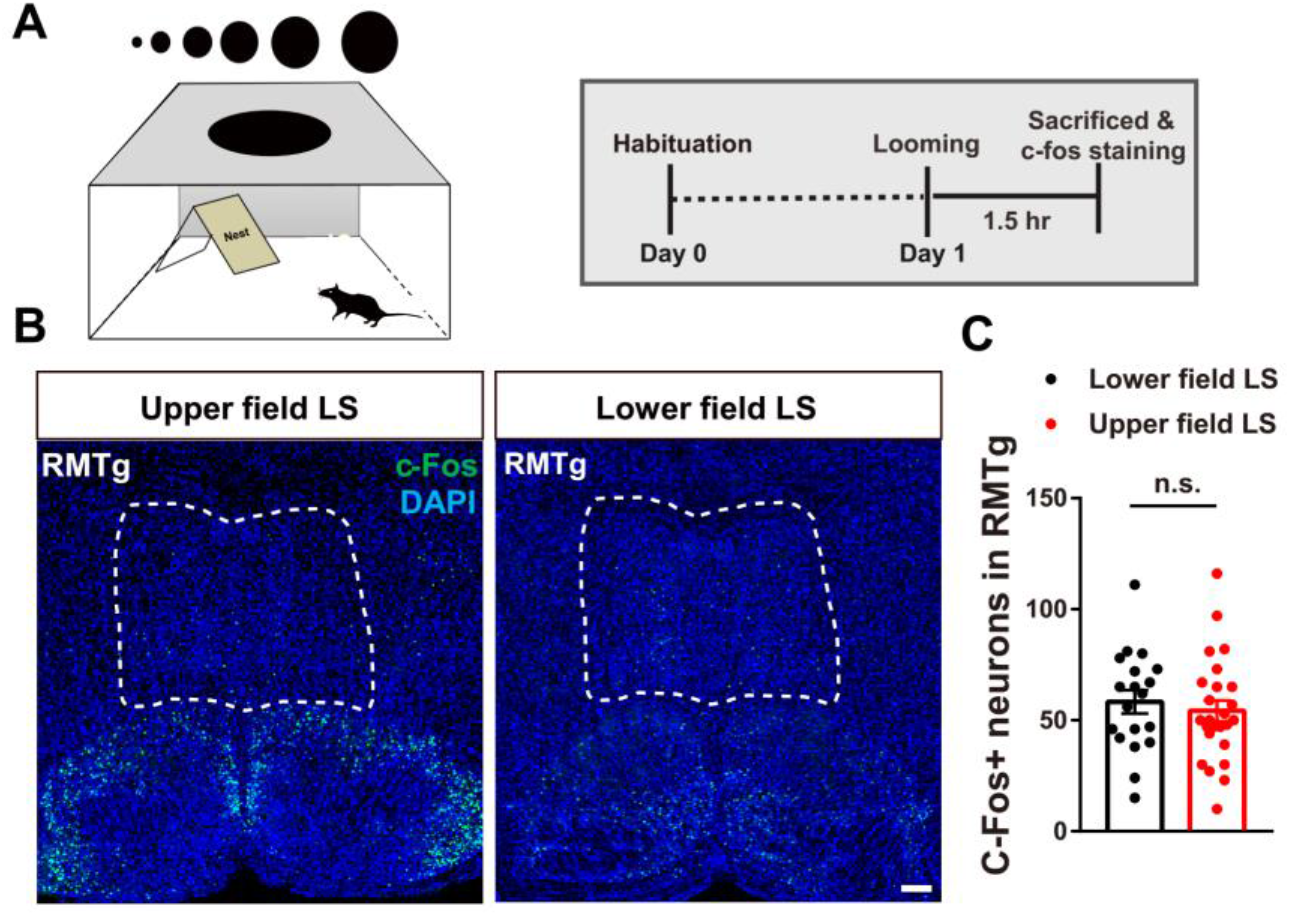
RMTg neurons did not respond to looming stimulus, which evokes defensive behavior. **(A)** Schematic paradigm of looming-evoked c-Fos expression. **(B)** Representative images of c-Fos expression in the VTA following upper field LS (left) and lower field LS (right) control stimulus (green, c-Fos; Blue, DAPI; scale bars, 100 μm,). **(C)** Upper field LS led to higher c-Fos expression in VTA TH negative neurons compared to lower field LS (n=19-25 slices from n_Lower field LS_ =4 mice, n_Upper field LS_ =6 mice, data presented as mean ± SEM, *t_42_* =0.5719, *P*=0.5704, Unpaired student test).

**Supplementary figure 2.**
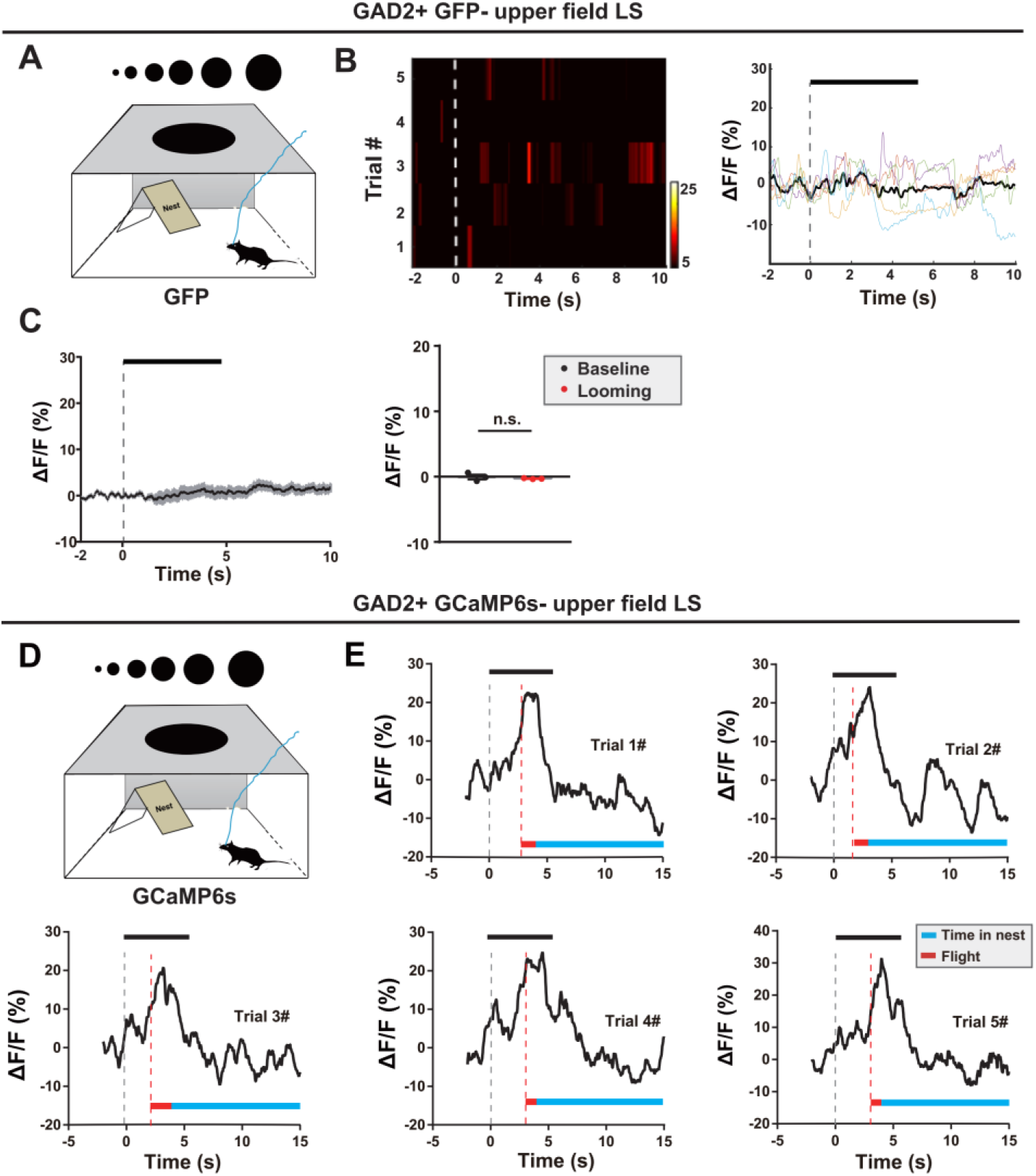
Upper field LS evoked significant increase of Ca2+ transients in VTA^GABA+^ neurons, but not in GFP group. **(A-B)** Representative trial-by-trial heatmap presentation of calcium transients, peri-event plot of the average calcium transient, and average calcium transients for the entire test group evoked by different visual stimuli, including upper field LS. **(C)** No change in calcium signal response following upper field LS was observed in GFP negative control mice (n = 3 mice; data presented as mean ± SEM, *t_2_*=0.6748, *P*= 0.5693, Paired student test). **(D-E)** Representative calcium transients signal from one mouse shows stable calcium signals across 5 trials of upper field LS and the signal’s relationship with upper field LS and the subsequent triggered flight to nest behavior. Gray dotted line indicates onset of looming; Red dotted line indicates onset of flight; Black bar represents the looming stimulus.

**Supplementary figure 3.**
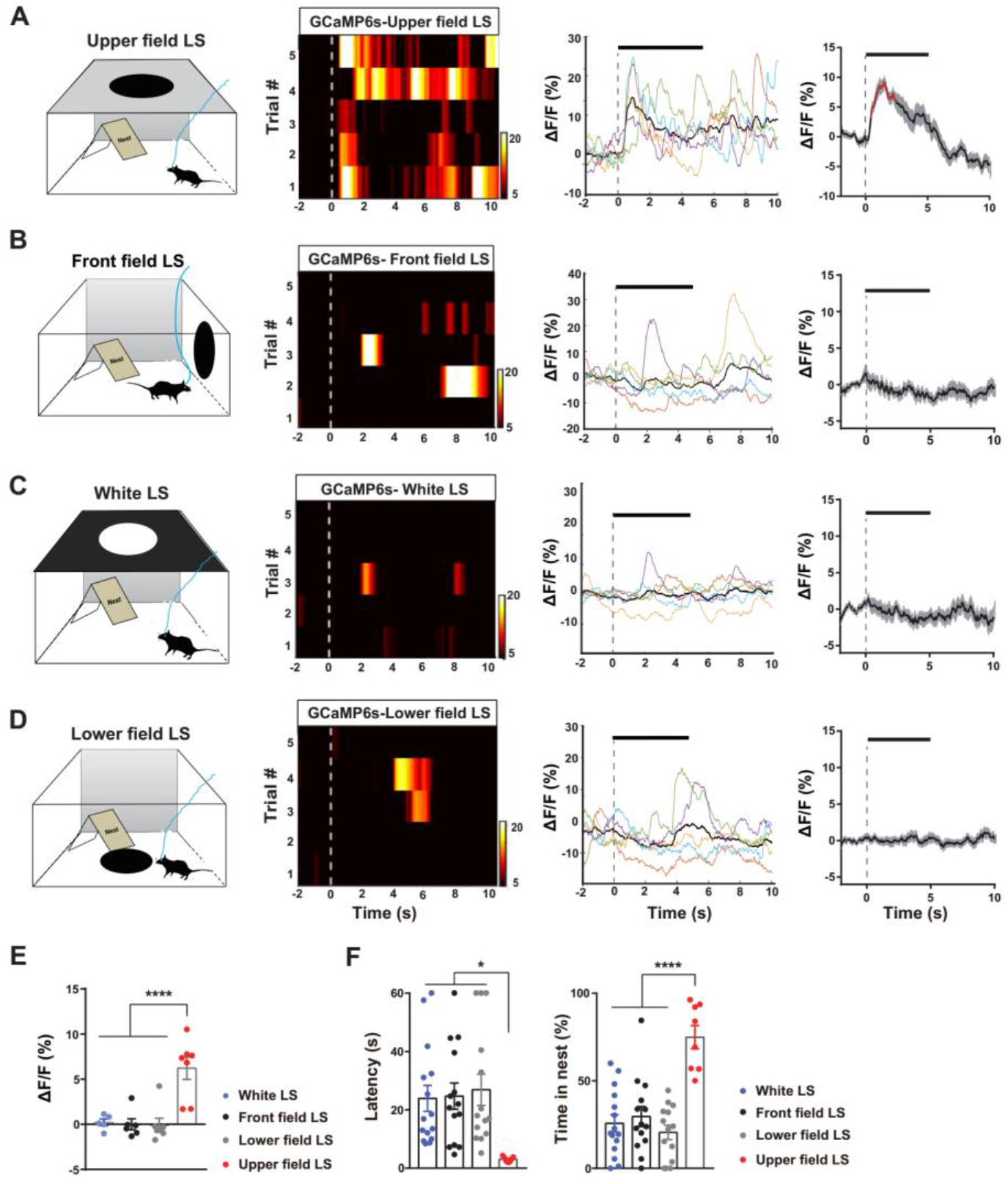
Upper field LS, but not other visual stimuli, evoked significant VTA^GABA+^ neuronal activation and flight-to-nest behavior. **(A-D)** Representative trial-by-trial heatmap presentation of calcium transients, peri-event plot of the average calcium transients, and average calcium transients for the entire test group evoked by different visual stimuli, including upper field LS (a), front field LS (b), white LS (c), and lower field LS (d). Red segments indicate statistically significant changes compared with the baseline (p<0.05; multivariate permutation tests. **(E)** Calcium transients were higher during upper LS than those during related visual stimuli, including front field LS, white LS and lower field LS (n_Front field LS_= 6 mice, n_Lower field LS_= 7 mice, n_white LS_= 5 mice, n_Upper field LS_= 7 mice; \*\*\*\**P*<0.0001, *F_3,21_* =13.17, one-way ANOVA). **(F)** Compared with the upper field LS, all the other visual stimuli including Collision, White LS and Lower field LS resulted in higher latency to return into nest after the stimulus and a shorter percentage of time spent in the nest after returning into the nest (n_Front field LS_ =14 mice; n_white LS_ =15 mice, n_Lower field LS_=14 mice; n_Upper field LS_=7 mice; \**P _latency_*= 0.0227, *F_3,46 latency_* = 3.501, \*\*\*\**P_time_* < 0.0001, *F_3,46 time_*= 13.01, one-way ANOVA). All data are presented as mean ± SEM.

**Supplementary figure 4.**
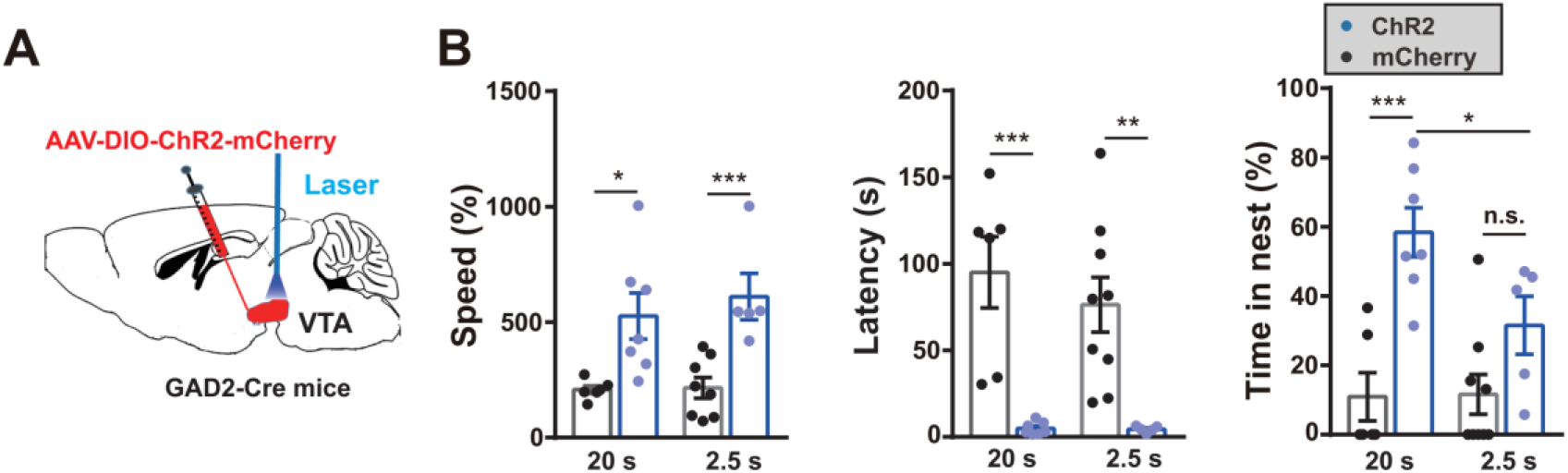
Longer activation of VTA^GABA+^ neurons elicited strong hiding time in nest. **(A)** Schematic diagram showing unilateral optogenetic activation of VTA GAD2+ neurons. **(B)** 2.5 s photoactivation of VTA GAD2+ neurons induced flight to nest behavior, shown by a decrease latency of flight-to-nest, an increase flight speed, and no change total percentage of hiding time in the nest than mCherry controls (n_mCherry 2.5s_= 8 mice, _n ChR2 2.5s_=5 mice, for latency, *t_11_*= 3.085, \**P*=0.0104; for speed, *t_11_*=4.089, \*\**P*=0.0018; for time in nest, *t_11_*=1.78, *P*=0.1028; Unpaired student test). 20 s photoactivation of VTA GAD2+ neurons induced flight to nest behavior, shown by a decrease latency of flight-to-nest, an increase flight speed, and increase percentage of hiding time in the nest than mCherry controls (n_mCherry 2.5s_= 6 mice, n_ChR2 2.5s_=7 mice, for latency, *t_11_*= 4.75, \*\*\**P*=0.0006; for speed, *t_11_*=2.899, \**P*=0.0145; for time in nest, *t_11_*=4.751, \*\*\**P*=0.0006; Unpaired student test). For the hiding time, 20s photoactivation induced longer hiding time in nest compared to 2.5 s (*t*_10_=2.457, \**P*=0.0338; Unpaired student test) All data are presented as mean ± SEM.

**Supplementary figure 5.**
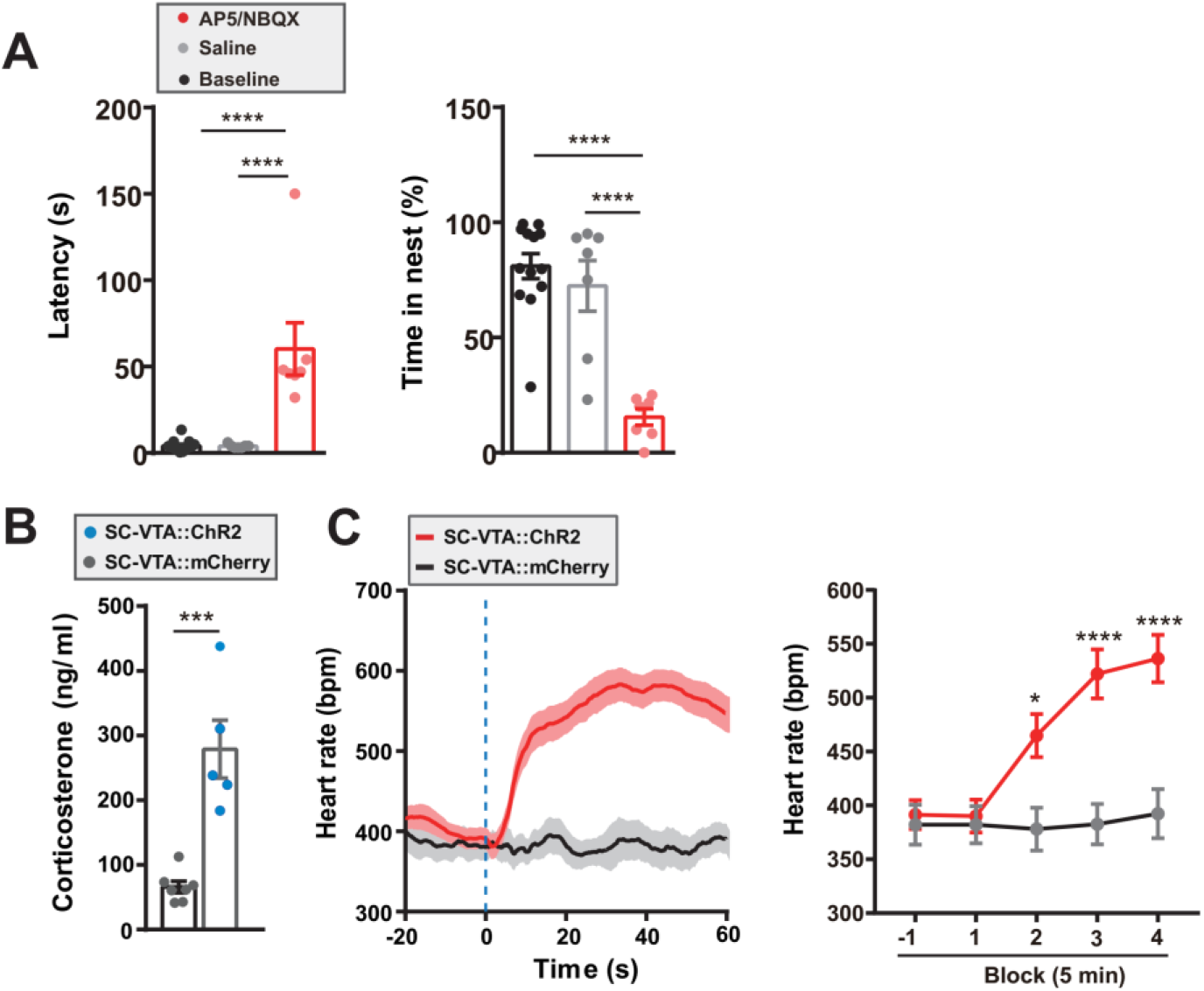
Activation of the SC–VTA pathway elicits defensive responses via glutamate receptor and accompanying increases in heart rate and circulating corticosterone levels. **(A)** Intra-VTA infusion of a glutamate antagonist AP5 and NBQX resulted in (left) higher latency to return to the nest and (right) a lower percentage of time spent in the nest after photostimulation compared to control groups (n_baseline_= 13 mice, n_saline_=7 mice, n_AP5+NBQX_=7 mice; \*\*\*\**P*<0.0001, *F_2, 24 time_*=20.1, *F_2, 24 latency_*= 24.15, bonferroni *post hoc* test, \*\*\*\**P*<0.0001, one-way *ANOVA test*). **(B)** Optical activation of CaMKIIα^SC-VTA^:: ChR2 led to higher mean plasma corticosterone levels than mCherry controls (n_mCherry_= 7 mice, n_ChR2_= 5 mice, *t_10_*=5.518, \*\*\**P*=0.0003; Unpaired student test). **(C)** Optical activation of CaMKIIα^SC-VTA^:: ChR2 resulted in higher mean heart rate than mCherry controls (n_mCherry_= 6 mice, n_ChR2_= 6 mice; Group x time effect interaction, \*\*\*\**P*< 0.0001, *F_4, 45_*= 6.009, two-way ANOVA with bonferroni *post hoc* analysis, \**P*=0.0486, \*\*\*\**P*< 0.0001. Each block represents 5 min). Blue dotted line represents onset of optical stimulation. All data are presented as mean ± SEM.

**Supplementary figure 6.**
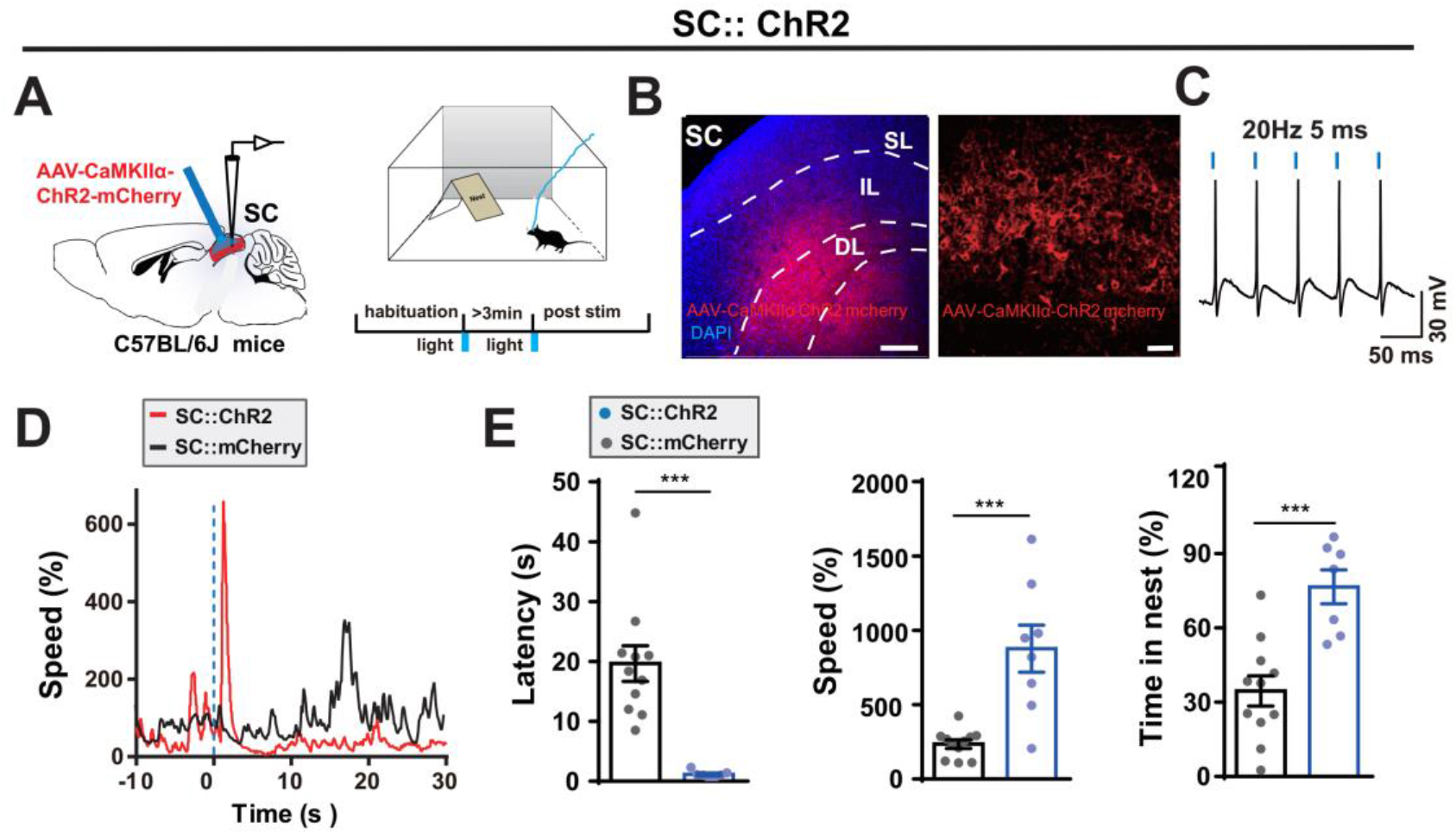
Optogenetic activation of the SC CaMKIIα-positive neurons evokes flight-to-nest behavior. **(A)** Schematic showing unilateral optogenetic activation of SC CaMKIIα-positive neurons in an open field with a nest apparatus. **(B)** Representative image showed ChR2 virus expression in the IL and DL layers of SC (blue, DAPI; red, ChR2-mCherry; scale bars, 250 μm, 50 μm respectively). **(C)** Example of light-induced action potentials from ChR2-mcherry+ neurons from SC using whole cellpatch-clamp slice recording. **(D)** Example of instant speed from two representative mice shows evoked flight behavior following opto-activation of ChR2-mouse compared with mCherry control (blue dotted line: onset of blue light optical stimulation). **(E)** Photoactivation of SC CaMKIIα-positive neurons induced flight to nest behavior of the animals, demonstrated by lower latency back into the nest, higher speed, and higher total percentage of time spent in the nest after looming stimulus compared to mCherry controls (n_mCherry_=11 mice, n_ChR2_= 7-8 mice, data were presented as mean ± SEM, for latency, *t_16_*=4.893, \*\*\**P*=0.0002; for speed, *t_17_*=4.668, \*\*\**P*=0.0002; for time in nest, *t_16_* = 4.452, \*\*\**P*=0.0004; Unpaired student test).

**Supplementary figure 7.**
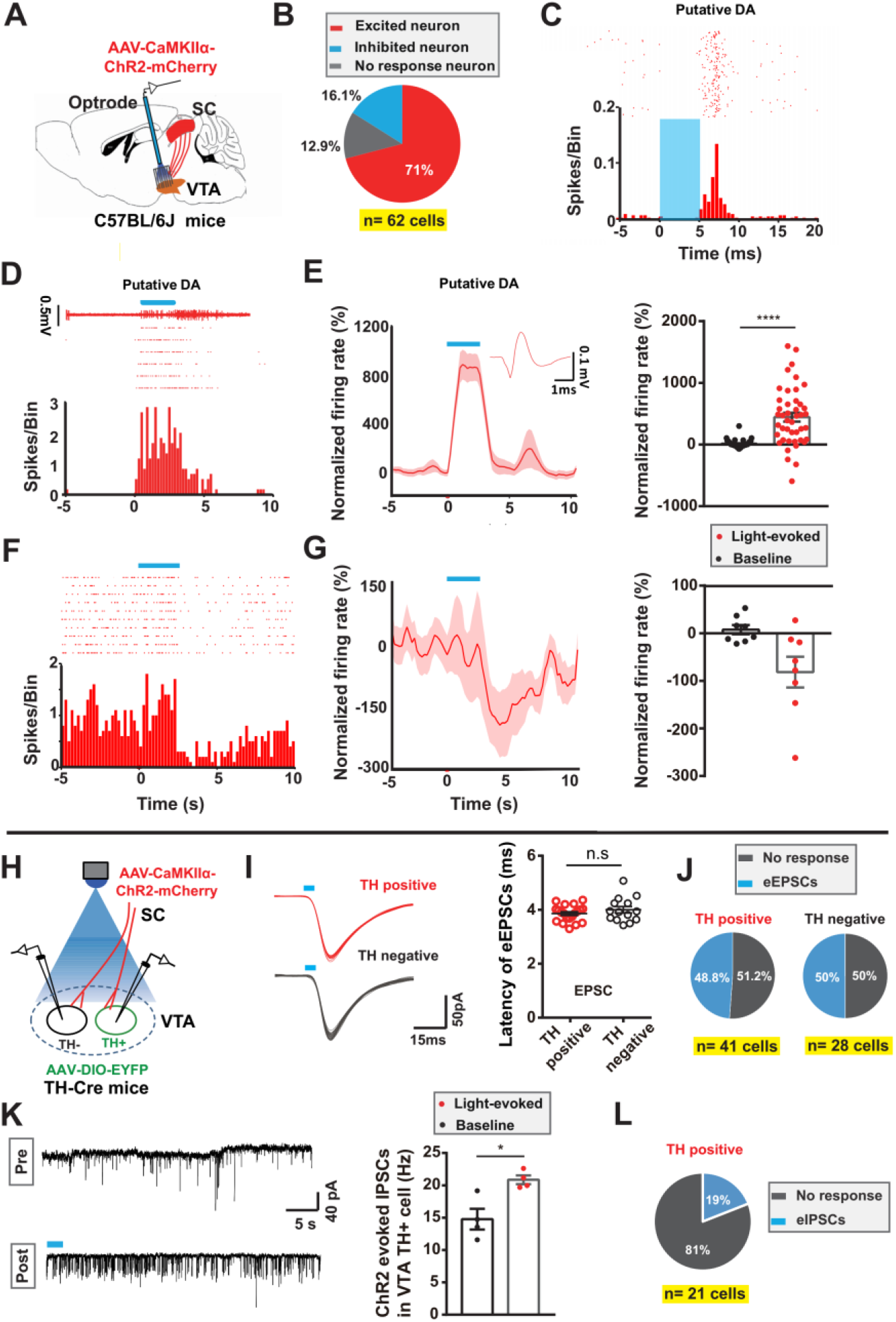
Activation of excitatory SC-VTA projections elicits VTA^DA+^ neuronal activation and inhibition. **(A)** *In vivo* multichannel recording of single-unit VTA neuron activity during optical stimulation of CaMKIIα SC-VTA terminals. **(B)** 44/62 putative DA neurons (71%) were excited, 8/62 (16.1%) were inhibited and 10/62 (12.9%) were unresponsive. **(C)** Representative PSTHs and raster plot of a single VTA putative DA neuron unit time-locked to 5 ms photostimulation. **(D)** Single-unit recording, PSTH and raster plot of a putative DA neuron excited by CaMKIIα SC-VTA terminals stimulation for 2.5 s. **(E)** Normalized firing rate of putative DA neurons shows the activation of VTA^DA+^ neurons by CaMKIIα SC-VTA fibers stimulation for 2.5 s (n= 44 cells; *t_43_*=5.86, \*\*\*\**P*< 0.0001, Paired student test). **(F)** Single-unit recording, PSTH and raster plot of a putative DA neuron inhibited by CaMKIIα SC-VTA fibers stimulation for 2.5 s. **(G)** Normalized firing rate of putative DA neurons shows the inhibition of VTA^DA+^ neurons by CaMKIIα SC-VTA fibers stimulation for 2.5 s (n= 8 cells; *t_7_* =2.631, \**P*= 0.0338, Paired student test). **(H)** Whole cell recording of eEPSCs in VTA TH+ and TH-neurons in brain slices during optogenetic stimulation of the CaMKIIα SC-VTA terminals. AAV-*EF1α*:: DIO-EYFP injections in TH-Cre mice were used to visualize TH+ neurons. **(I)** Left, example of eEPSCs in TH+ and TH-neurons in the VTA evoked by CaMKIIα SC-VTA terminals stimulation; Right, no difference in latency of the eEPSCs between VTA TH+ and TH-neurons (n_TH+_= 20 cells from 17 mice; n_TH-_=14 cells from 12 mice, latency _TH+_= 3.86 ms± 0.07, latency _TH-_=4.00 ms± 0.12, *P*= 0.2814, *t_32_*=1.096, unpaired student test); **(J)** Left, of 41 TH+ neurons, 51.2% neurons were non-responsive while 48.8% evoked EPSCs following CaMKIIα SC-VTA terminals stimulation (n total=41 cells). Right, of 28 TH-neurons, 50% neurons were non-responsive while 50 % evoked EPSCs following CaMKIIα SC-VTA terminals stimulation (n_total_=28 cells). **(K)** Left, example of increase of light evoked inhibitory postsynaptic currents (eIPSCs) frequency from TH+ neurons from patch-clamp slice recording; Right, eIPSCs were frequency induced by CaMKIIα SC-VTA terminals stimulation (n=4 cells from 4 mice, *t_3_*= 4.213, \**P*= 0.0244, Paired student test). **(L)** Of 21 TH+ neurons, 81 % were non-responsive while 19% evoked IPSCs following CaMKIIα SC-VTA terminals stimulation (n_total_=21 cells). All data are presented as mean ± SEM.

**Supplementary figure 8.**
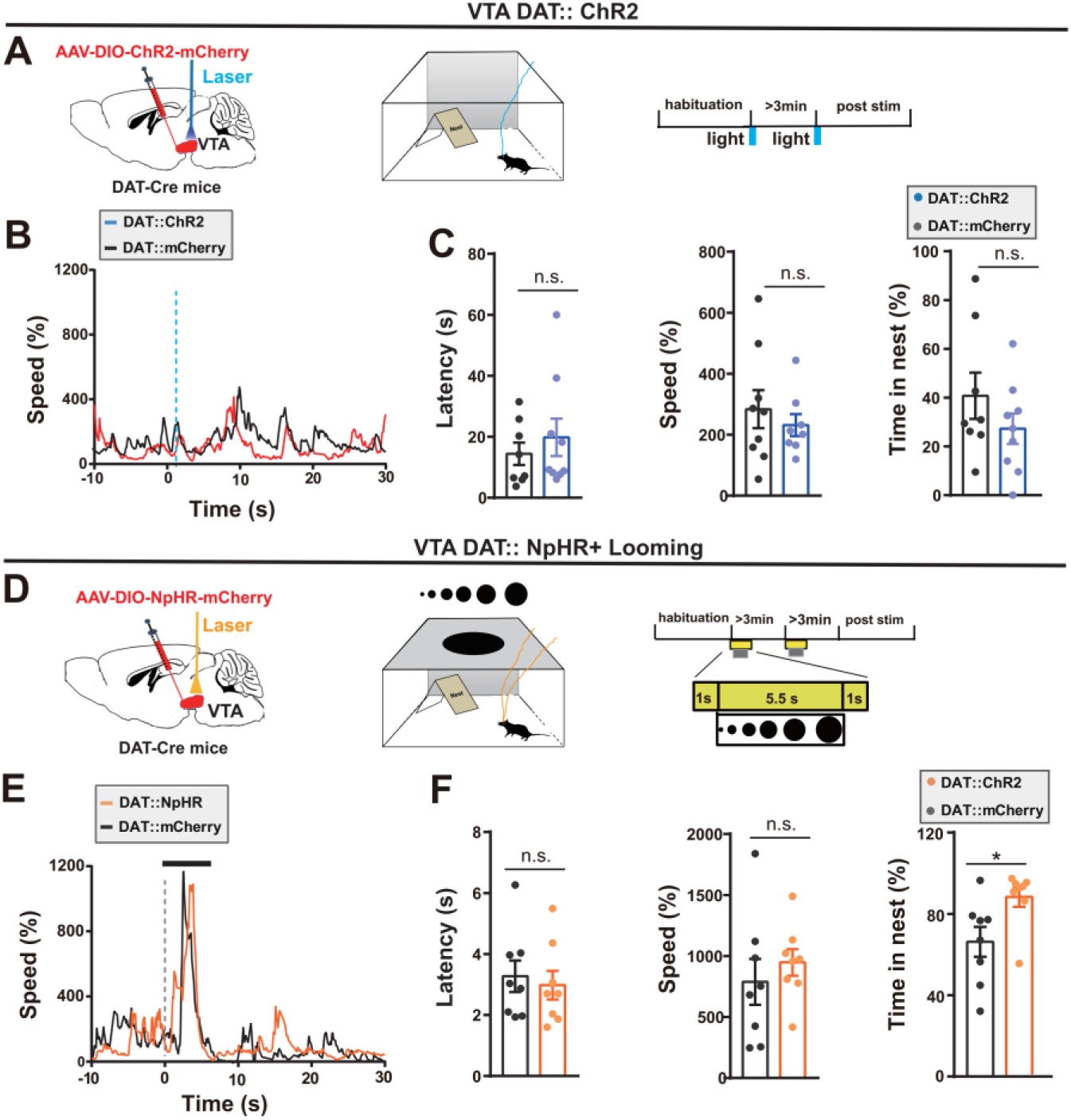
VTA^DA+^ neurons do not mediate looming-evoked flight-to-nest behavior. **(A)** Optogenetic activation of VTA^DA+^ neurons using DAT-Cre mouse during upper field looming stimulus in an open field with a nest. **(B)** Example of instant speeds from a representative mouse from each group evoked by blue light (blue dotted line: onset of optical stimulation). **(C)** Photostimulation of VTA DAT+ neurons did not result in any change in the speed, latency back to the nest, or total percentage of time spent in the nest following looming-evoked flight to the nest behavior (n_mCherry_= 8 mice, n_ChR2_= 9 mice, for latency, *t_15_*= 0.7364, *P*=0.4728; for speed, *t_15_*=0.6985, *P*=0.4955; for time in nest, *t_15_*= 1,211, *P*=0.2447; Unpaired student test). **(D)** Optogenetic bilateral inhibition of VTA^DA+^ neurons using DAT-Cre mouse during upper field looming stimulus in an open field with a nest. **(E)** Example of instant speeds from a representative mouse from each group illustrates flight behavior evoked by looming (gray dotted line: onset of looming stimulation). **(F)** Photoinhibition of VTA DAT+ neurons result in no change in the speed or latency back to the nest but increased total percentage of time spent in the nest following looming-evoked flight to the nest behavior (n_mCherry_= 8 mice, n_NpHR_= 8 mice, for latency, *t_14_*= 0.4154, *P*=0.6841; for speed, *t_14_*=0.7358, *P*=0.4740; for time in nest, *t_14_*=2.513, \**P*=0.0248; Unpaired student test). All data are presented as mean ± SEM.

**Supplementary figure 9.**
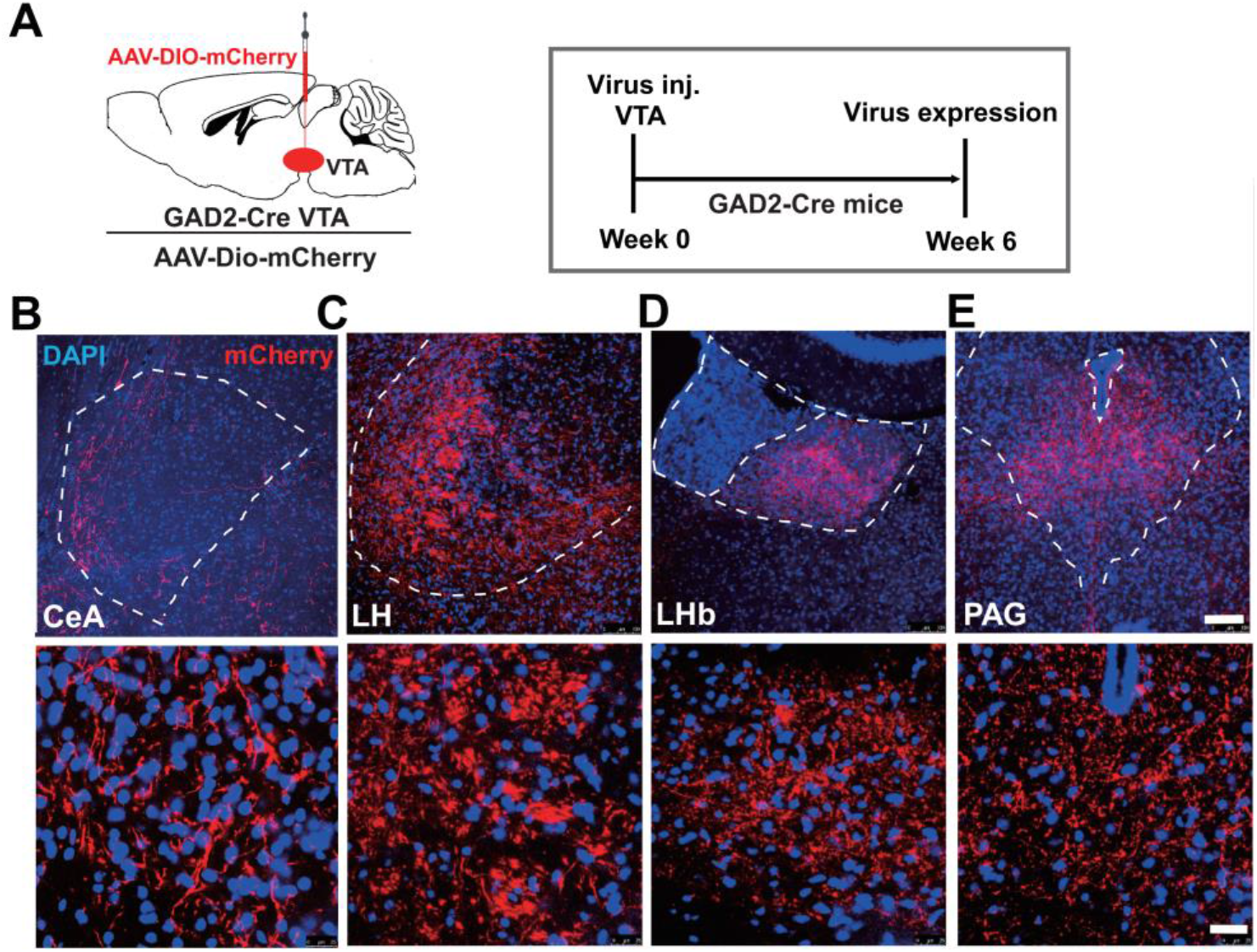
Screening of VTA^GABA+^ neurons output regions. **(A)** AAV-*EF1α*:: DIO-mCherry injection into the VTA of GAD2-cre mice. **(B)** Anterograde tracing of VTA^GABA+^ neurons shows fibers in the CeA (b), LH (c), LHb (d), and PAG (e) with high-magnification images of the terminal fibers (scale bars, 100 μm and 20 μm respectively).

**Supplementary figure 10.**
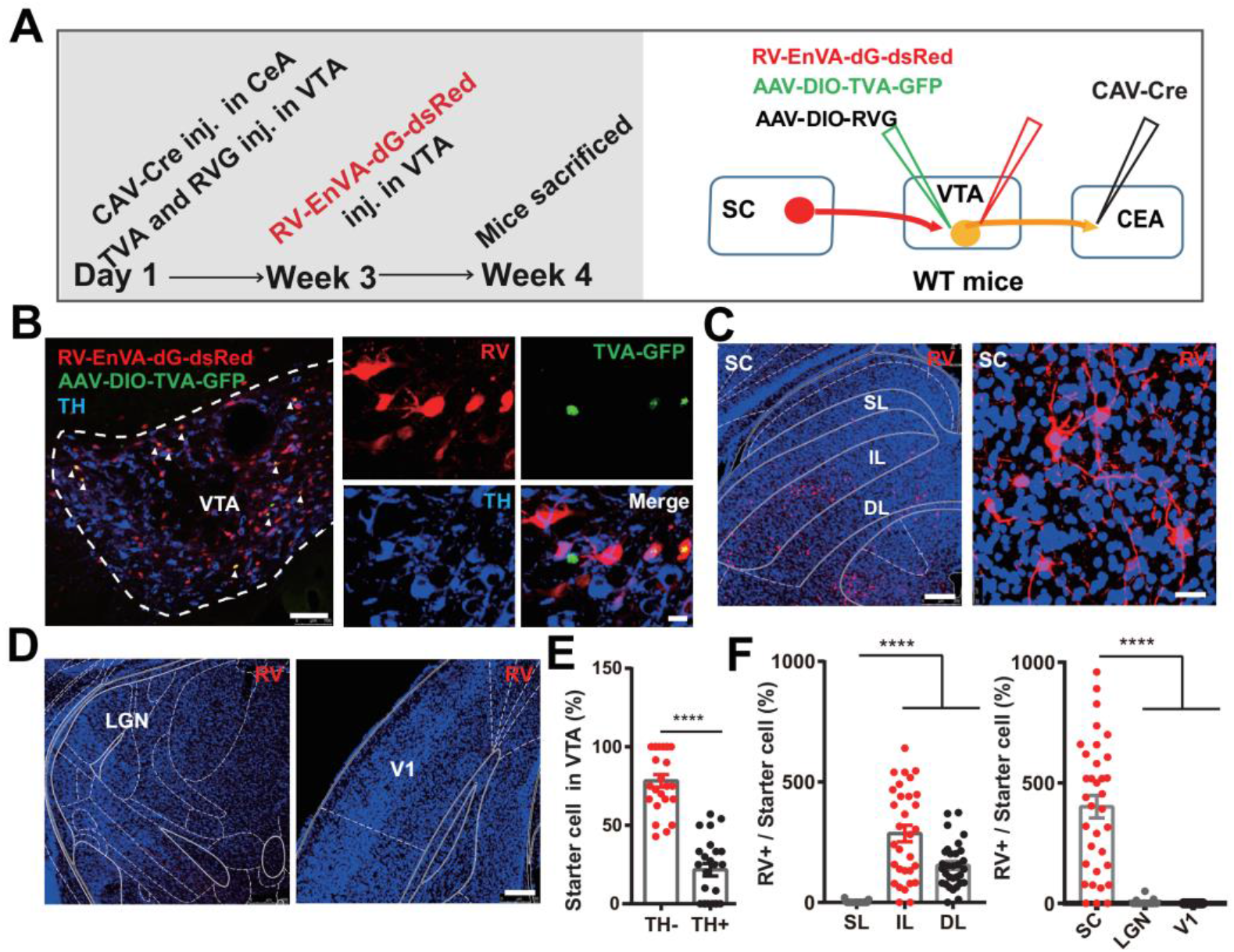
TRIO tracing identifying the input-output relationship of VTA neurons. **(A)** Trace the input-output relationships of VTA neurons: CAV-Cre virus was injected into the CeA. Rabies-EnVA-dG-dsRed, AAV-*EF1α*:: DIO-Rabies-G and AAV-*EF1α*:: DIO-TVA-GFP was co-injected into the VTA. **(B)** Representative image showing the co-expression of rabies-EnVA-dG-dsRed (red), AAV-*EF1α*:: DIO-RVG, and AAV-*EF1α*:: DIO-TVA-GFP (green) in the VTA with high-magnification images of VTA starter cells (yellow) (scale bars, 250 μm and 10 μm respectively). White arrows indicate the starter cells in VTA. **(C-D)** Representative images showing substantial rabies-dsRed signal from CeA-projecting VTA neurons in the IL and DL layers of SC, whilst only rare signals were observed in other visual related brain regions, LGN and V1 (blue, DAPI, red, rabies-dsRed; scale bar, 250 μm and 20 μm respectively). **(A)** Quantification analysis showed that 78.36 ± 3.99% of the CeA-projecting VTA starter cells were TH negative (n=24 slices from 3 mice, data presented as mean ± SEM, *t_46_*=10.05, \*\*\*\**P*<0.0001; Unpaired student test). **(B)** Left, quantification of the number of rabies-dsRed labeled neurons in SC subregions: intermedial layer (IL), deep layer of the SC (DL) and superficial layer (SL); (n =31 slices from 4 mice, data presented as mean ± SEM, \*\*\*\**P*<0.0001, *F_2, 90_*= 57.29; one-way ANOVA). Right, quantification of the number of rabies-dsRed labeled neurons in the SC, V1, and LGN (n=23-31 slices from 3-4 mice, data presented as mean ± SEM, \*\*\*\**P*<0.0001, *F_2, 79_*=59.45, one-way ANOVA)

**Supplementary figure 11.**
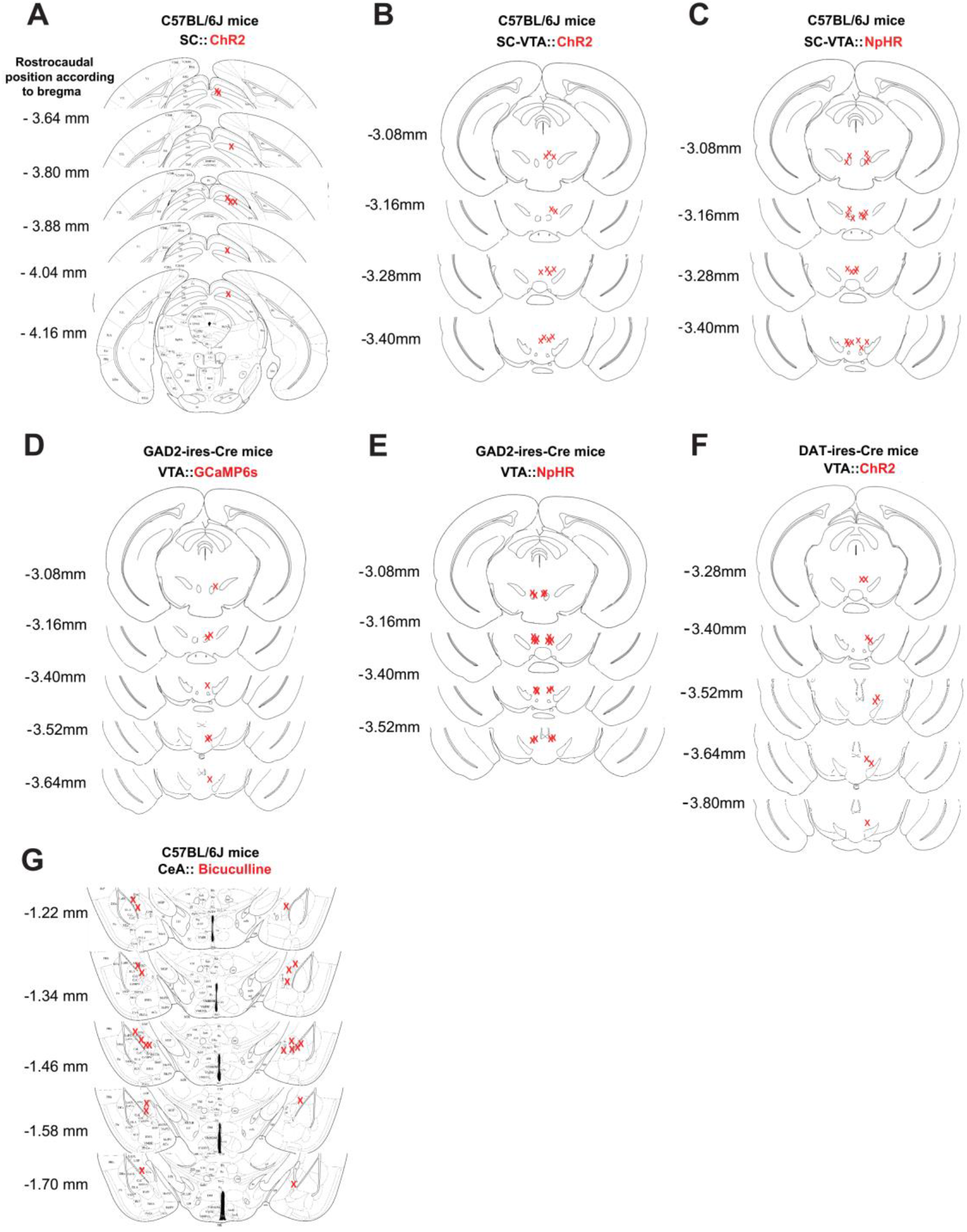
Optical fiber placements of optogenetic stimulation, fiber photometry and cannula placements of pharmacology. Location of optical fibers within SC and VTA **(A-F)**. Location of cannula within CeA **(G)**. All placements based on histological staining of brain slices after experiments.

## STAR METHODS KEY RESOURCES TABLE

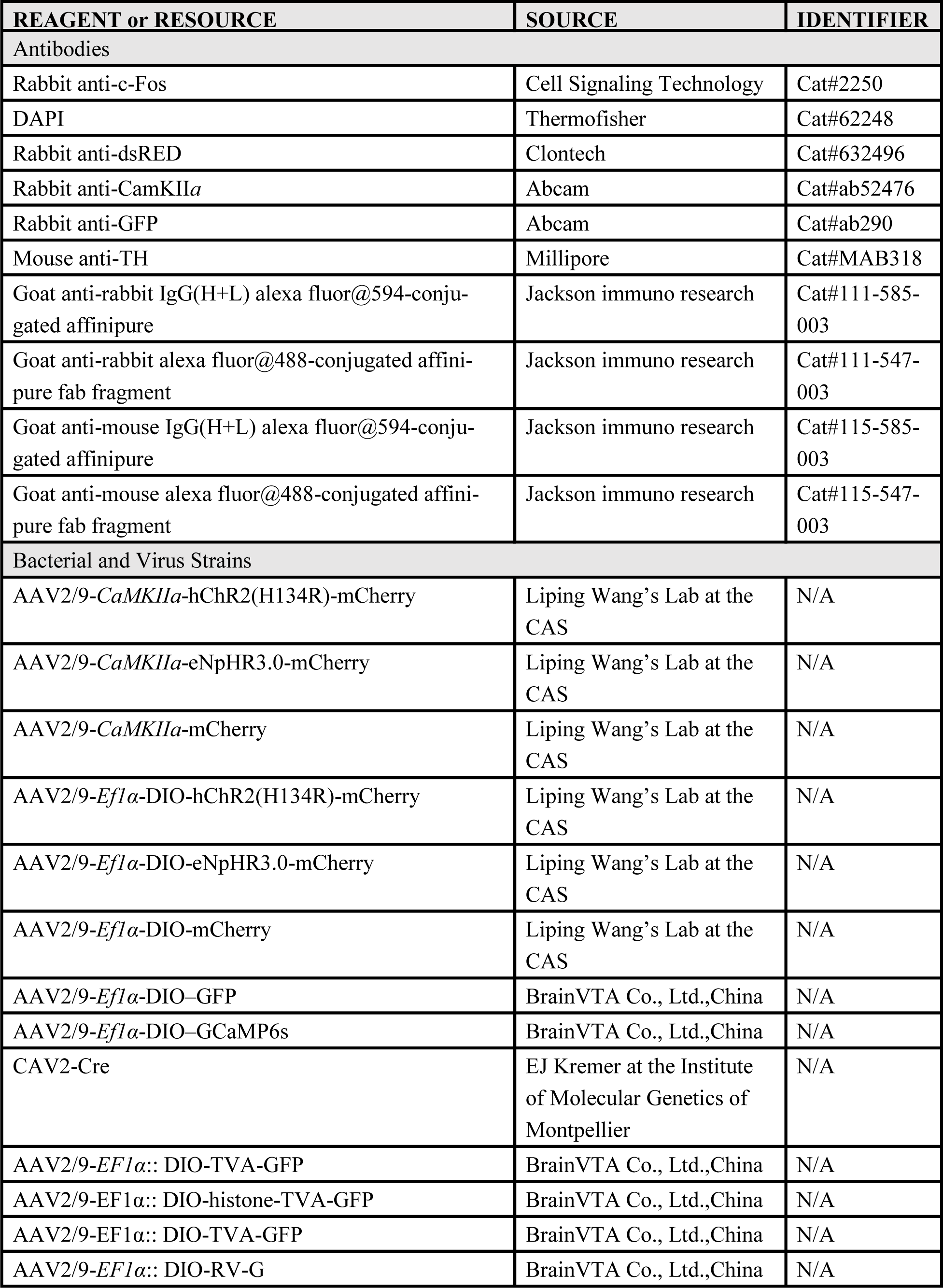
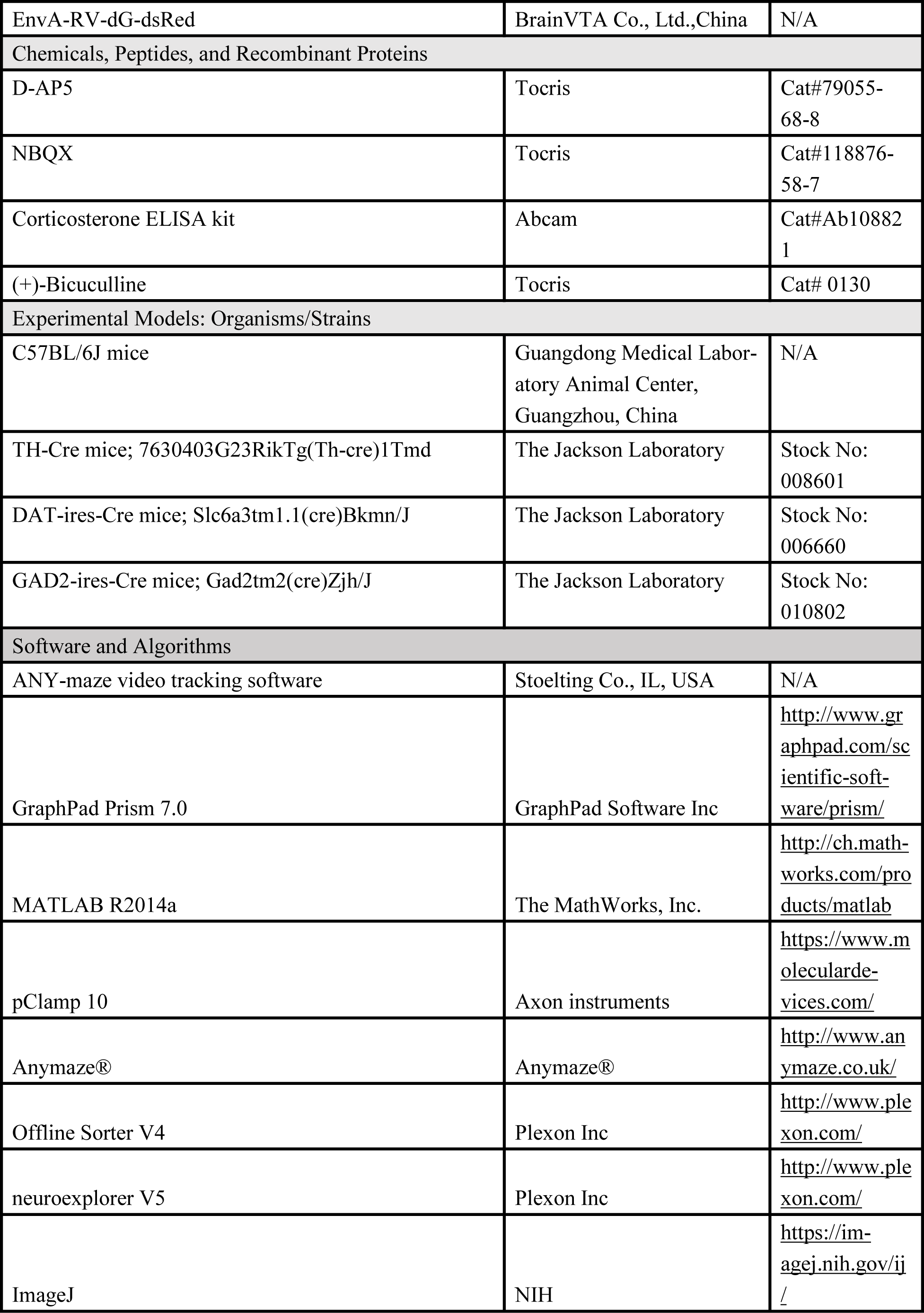
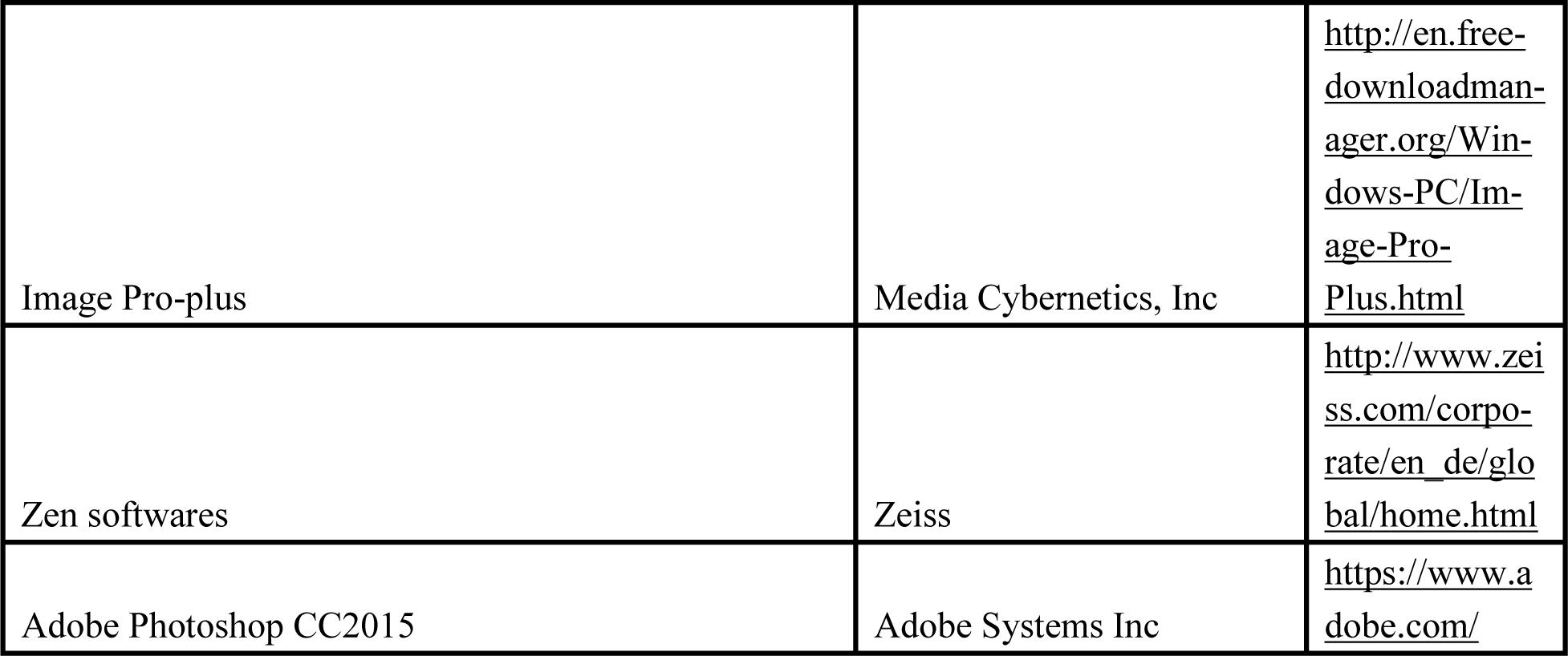

## CONTACT FOR REAGENT AND RESOURCE SHARING

Further information and requests for resources and reagents should be directed to and will be fulfilled by the Lead Contact, Liping Wang; Fuqiang Xu

## EXPERIMENTAL MODEL AND SUBJECT DETAILS

### Animals

All husbandry and experimental procedures in this study were approved by the Animal Care and Use Committees at the Shenzhen Institute of Advanced Technology (SIAT) or Wuhan Institute of Physics and Mathematics (WIPM), Chinese Academy of Sciences (CAS). Adult (6 to 8 week-old) male C57BL/6 (Guangdong Medical Laboratory Animal Center, Guangzhou, China), TH-Cre (Jax No. 008601) (Savitt, 2005), GAD2-ires-Cre (Jax No. 010802) (Taniguchi et al., 2011) and DAT-ires-Cre (Jax No. 006660) (Ekstrand et al., 2007) mice were used in this study. Mice were housed at 22–25 °C on a circadian cycle of 12-hour light and 12-hour dark with ad-libitum access to food and water.

## METHOD DETAILS

### Viral vector preparation

For optogenetic experiments, we used plasmids for AAV2/9 viruses encoding *CaMKIIα*:: hChR2 (H134R)–mCherry, *CaMKIIα*:: eNpHR3.0–mCherry, *CaMKIIα*:: mCherry, *EF1α*:: DIO–hChR2 (H134R)–mCherry, and *EF1α*:: DIO–eNpHR3.0–mCherry (all gifts from Dr. Karl Deisseroth, Stanford University). Viral vector titers were in the range of 3-6×10^12^ genome copies per ml (gc)/mL. For AAV2/9 viruses encoding *EF1α*:: DIO–GCaMP6s and *EF1α*:: DIO–GFP were all packaged by BrainVTA Co., Ltd., Wuhan. For rabies tracing, viral vectors AAV2/9-*EF1α*-:: DIO-TVA-GFP, AAV2/9-*EF1α*:: DIO-histone-TVA-GFP, AAV2/9-*EF1α*:: DIO-TVA-GFP, AAV2/9-*EF1α*:: DIO-RV-G, and EnvA-RV-dG-dsRed were all packaged by BrainVTA Co., Ltd., Wuhan. Adeno-associated and rabies viruses were purified and concentrated to titers at approximately 3×10^12^ v.g/ml and 1×10^9^ pfu/ml, respectively. The canine adenovirus type-2 encoding Cre recombinase (Kremer et al., 2000) (CAV2-Cre, 4×10^12^ v.g/ml) was provided by EJ Kremer at the Institute of Molecular Genetics of Montpellier.

### Virus injection

Animals were anesthetized with pentobarbital (i.p., 80 mg/kg), and then placed in a stereotaxic apparatus (RWD, China). During surgery and virus injections, animals were kept anesthetized with isoflurane (1%). The skull above targeted areas was thinned with a dental drill and carefully removed. Injections were conducted with a 10 μl syringe connected to a 33-Ga needle (Neuros; Hamilton, Reno, USA), using a microsyringe pump (UMP3/Micro4, USA). Experiments were performed at least 5-8 weeks after virus injection. Fiber implantation coordinates for optical stimulation of the SC were: anterior posterior (AP), −3.80 mm; medial lateral (ML), ±0.80 mm; dorsoventral (DV), −1.8 mm. VTA coordinates were: AP −3.20 mm, ML ±0.25 mm, and DV −4.4 mm. Amygdala coordinates were: AP, −1.5mm; ML, ±2.95 mm; DV, −4.75 mm. Viruses were delivered unilaterally for ChR2 and bilaterally for eNpHR3.0, GCaMP6s and GFP.

### Trans-synaptic tracer labeling

All animal procedures were performed in Biosafety level 2 (BSL2) animal facilities. To determine whether the SC-VTA pathway was innervated by GABAergic neurons in the VTA, Gad2-Cre mice (20-25 g) were used for trans-mono-synaptic tracing based on the modified rabies virus. A mixture of AAV2/9-*EF1α*:: DIO-TVA-GFP and AAV2/9-*EF1α*::DIO-RV-G (1:1, total volume of 150 nl) was injected into the VTA region using the following coordinates: AP, −3.20 mm; ML, −0.25 mm; DV, −4.40 mm. Three weeks later, 200 nl of EnvA-pseudotyped rabies virus (EnvA-RV-dG-DsRed) was injected into the VTA using the previously defined coordinates.

To identify the SC-VTA-CeA neural pathway, we performed a *tracing the relationship between input and output* (TRIO) experiment. On the first day, we injected 150 nl of CAV2-Cre into the CeA (AP, −1.50 mm; ML, 2.95 mm; DV, −4.75 mm) of adult male C57BL/6 mice. On the same day, we injected a mixture of 150 nl AAV2/9-*EF1α*:: DIO-histone-TVA-GFP and AAV2/9-*EF1α*:: DIO-RV-G (1:1) into the VTA (AP, −3.20 mm; ML, −0.25 mm; DV, −4.40 mm) of these animals. Three weeks later, we injected 200 nl of EnvA-RV-dG-DsRed into the VTA using the same coordinates as before. Mice were sacrificed one week after RV injection.

To determine whether the SC-VTA-CeA pathway was innervated by GABAergic neurons in the VTA, we modified the mono-synaptic rabies tracing strategy. On the first day, we injected a mixture of 150 nl AAV2/9-*EF1α*:: DIO-histone-TVA-GFP and AAV2/9-*EF1α*:: DIO-RV-G (1:1) into the VTA of GAD2-ires-Cre mice. Six weeks later, when the accumulated TVA of GABAergic VTA neurons was transported to axon terminals in the CeA, we injected 200 nl of EnvA-RV-dG-dsRed into the CeA (AP, −1.5 mm; ML, 2.95 mm; DV, −4.75 mm) of these mice. Thus, we specifically infected CeA-projecting GABAergic VTA neurons and traced their inputs. Mice were sacrificed one week after RV injection.

### Implantation of optical fiber(s) and cannulas

To optically stimulate terminals, a 200 μm optic fiber (NA: 0.37; NEWDOON, Hangzhou) was unilaterally implanted into the SC (AP, −3.8 mm; ML, −0.6 mm; DV, −1.4 mm) and VTA (AP, −3.20 mm; ML, −0.25 mm; DV, −3.8 mm). For the inhibition of neuron soma or projections, optic fibers were bilaterally implanted into the VTA (AP, −3.20 mm; ML, ±1.50 mm; DV, −4.0 mm) at a 15° angle from the vertical axis. For fiber photometry, the optic fiber was bilaterally implanted into the VTA of GAD2-Cre mice (AP, −3.20 mm; ML, −1.5 mm; DV, −4.5 mm) at a 15° angle. For pharmacological experiments, drug cannulas were bilaterally implanted into the VTA (AP, −3.20 mm; ML, ±0.35 mm; DV, −3.8 mm) and CeA (AP, −1.3 mm; ML, ±2.95 mm; DV, −4.2 mm). Mice had at least 2 weeks to recover after surgery.

### Patch-clamp electrophysiology

We use standard procedures to prepare coronal slices (300 μm) from 14-16 weeks old GAD2-ires-Cre and TH-Cre mice, which had received virus injections six weeks earlier. Recordings in VTA were made on visually identified neurons expressing EYFP. Coronal sections were cut with a vibratome (Leica) into a chilled slicing solution containing the following (in mM): 110 Choline Chloride, 2.5 KCl, 1.3 1.3 NaH_2_PO_4_, 25 NaHCO_3_, 1.3 Na-Ascorbate, 0.6 Na-Pyruvate, 0.5 CaCl_2_, 7 MgCl_2_). Then, slices were incubated at 32 °C for 30 min in artificial cerebrospinal fluid (ACSF) which contained (in mM): 125 NaCl, 2.5 KCl, 1.3 NaH_2_PO_4_, 25 NaHCO_3_, 1.3 Na-Ascorbate, 0.6 Na-Pyruvate, 10 Glucose, 2 CaCl_2_, 1.3 MgCl_2_ (pH 7.35 when saturated with 95% O_2_/ 5% CO_2_), and allowed to equilibrate to room temperature for >30 min. The osmolarity of all solutions was maintained at 280–300 mOsm.

Evoked EPSCs were induced using 5 ms blue light stimulation of VTA terminals of CaMKIIα positive neurons expressing ChR2 from SC soma projecting to VTA. Recordings were performed on GAD2+ neurons (EYFP positive), TH+ (EYFP positive) or TH-(EYFP negative) with pipettes filled with the following (in mM): (105 Potassium gluconate, 30 KCl, 10 HEPES, 10 phosphocreatine, 0.3 EGTA, 5 QX314, 4 Mg-GTP, 0.3 Na-ATP, pH 7.35. To identify the eEPSCs glutamatergic nature, ionotropic glutamate receptor antagonists, d-2-amino-5-phosphonovalerate (AP-5; 25 μM) and 2, 3-dihydroxy-6-nitro-7-sulfamoyl-benzoquinoxaline-2, 3-dione (NBQX; 20 μM) were added at the end of recordings.

Evoked IPSCs were elicited using blue light (5 ms pulse, 60Hz) stimulation of CeA axon terminals of GABAergic neurons expressing ChR2 VTA axons projecting to CeA. Recordings were performed on CeA neurons with pipettes filled with the following (in mM): 130 mM cesium gluconate, 7 mM CsCl, 10 mM HEPES, 2 mM MgCl_2_, 4 mM Mg-ATP, 0.3 mM Tris-GTP, and 8 mM QX314, pH 7.25. To rule out glutamatergic inputs, ionotropic glutamate receptor antagonists, AP-5 (25 μM) and NBQX (20μM) were add to the artificial cerebrospinal fluid. To identify the eIPSCs GABAergic nature, GABA-A receptors with bicuculline (25 μM) was added at the end of recordings.

Several criteria had to be met for successful inclusion of recording data: 1) The amplitude of eIPSCs or eEPSCs was higher than 10 pA; 2) latency was less than 10 ms for at least 60% of the trials. 3) Whole-cell patch-clamp recordings were discarded if the access resistance exceeded 10 MΩ and changed more than 25% during the recordings.

Pipettes were formed by a micropipette puller (Sutter P-2000) with a resistance of 3–5 MΩ. During whole-cell patch recording, we viewed individual cells with an up-right fixed-stage microscope (FN-S2N; Nikon., Japan) equipped with a water immersion objective (40×, 0.8 numerical aperture), IR-filtered light, differential interference contrast optics, and a Coolsnap HQ CCD camera (Photomatrics, Britannia). All recordings were conducted with a MultiClamp700B amplifier (Molecular Devices). Analog signals were low-pass filtered at 2 kHz, digitized at 20 kHz using Digidata 1440A, and recorded using pClamp 10 software (Molecular Devices).

Data are presented as means ± standard error of the mean (SEM). Statistical significance was determined using a two-tailed Student’s t-test or a two-way analysis of variance (ANOVA), with a significance level of P<0.05.

### Histology, immunohistochemistry, and microscopy

Mice received an overdose of chloral hydrate (10% W/V, 300 mg/kg body weight, i.p.) and were then transcardially perfused with cold phosphate-buffered saline (PBS), followed by ice-cold 4% paraformaldehyde (PFA; Sigma) in PBS. Brains were removed and submerged in 4% PFA at 4 °C overnight to post-fix, and then transferred to 30% sucrose to equilibrate. Coronal brains sections (40 μm) were obtained on a cryostat microtome (Lecia CM1950, Germany). Freely floating sections were washed with PBS, blocking solution (0.3% TritonX-100 and 10% normal goat serum, NGS in PBS, 1 h at room temperature). Sections were then incubated in primary antiserum (rabbit anti-c-Fos, 1:300, Cell Signaling; Rabbit anti-CaMKIIα, 1:250, Abcam; 1:1000, Abcam; rabbit anti-TH, 1:500, Abcam; mouse anti-TH, 1:500, Millipore; rabbit anti-dsRed,1:1000, Clontech; rabbit anti-GFP, 1:500, Abcam) diluted in PBS with 3% NGS and 0.1% TritonX-100 overnight. The secondary antibodies used were Alexa fluor 488, 594, or 405 goat anti-mouse IgG and Alexa fluor 488, 594, or 405 goat antirabbit (all 1:200, Jackson) at room temperature for 1 h. Sections were mounted and cover slipped with anti-fade reagent with DAPI (ProLong Gold Antifade Reagent with DAPI, life technologies) or signal enhancer (Image-iT FX Signal Enhancer, Invitrogen). Sections were then photographed and analyzed with a Leica TCS SP5 laser scanning confocal microscope and ImageJ, Image Pro-plus, and Photoshop software.

For the looming-evoked c-Fos staining experiment, mice were sacrificed 1.5 hr post looming stimulus and brains then subjected to c-Fos staining. The imagines were taken and then overlaid with The Mouse Brain in Stereotaxic Coordinates to locate the VTA (with coordinates from bregma: −2.9~3.8 mm). Then the c-Fos staining was manually counted by an individual experimenter blind to the experiment groups.

### Plasma corticosterone measurement

Animals were euthanized by rapid decapitation and trunk blood was collected into heparinized tubes 10 min after optic stimulation (50 pulses, 20 Hz blue light, 5 ms pulse duration). After centrifugation of the blood at 3000 rpm for 20 min at 4 °C, the serum was stored at −80 °C until assay. Plasma corticosterone level was measured using a commercially available ELISA kit (Abcam).

### Electrocardiogram recording

Heart rate (HR) recordings were measured using the MouseOX® Plus non-invasive pulse oximeter (STARR Life Sciences, Oakmont, PA). The neck collar and system was set up according to manufacturer instructions. After a baseline recording of 5 min, 5 photostimulation trials were applied (50 pulses, 20 Hz, 3 min inter-spike interval [ISI]). Data were extracted using WINDAQ software (© DATAQ). HR data was then analyzed with custom-written Matlab scripts. HR time courses were obtained by averaging data from each trial around the time of stimulation. Error bars represent mean ± SEM.

### Looming test

The looming test was performed in a 40 x 40 x 30 cm closed Plexiglas box with a shelter nest in the corner. For upper field LS, an LCD monitor was placed on the ceiling to present multiple looming stimulus. For upper visual field LS, an LCD monitor was placed on the ceiling to present multiple looming stimuli, which was a black disc expanding from a visual angle of 2° to 20° in 0.3 s, i.e., expanding speed of 60°/s. The expanding disc stimulus was repeated for 15 times in quick succession (totally 4.5 s). This together with a 0.066 s pause between each repeat, the total upper visual field LS last 5.5 s.

Lower field LS (same stimulus as above but presented to the lower field), upper field white LS (a disc of reversed contrast (white on gray) presented to the upper field) and front field LS (the same black stimulus except presented to the front field) were used as control visual stimulus.

Behavior was recorded using an HD digital camera (Sony, Shanghai, China). Animals were handled and habituated for 10-15 min to the looming box one day before testing. During the looming test session, mice were first allowed to freely explore the looming box for 3-5 min. For optogenetic or pharmacological experiment plus looming experiment, we performed 2 trials of looming stimuli while only the first defensive behavior output was analyzed; for c-Fos and calcium signal experiment, total 5 trials of looming stimuli were presented and analyzed. No observable adaptation was observed in all our experiments.

The optogenetic inhibition/stimulation and the looming stimulus were coupled by ARBITRARY/FUNCTION GENERATOR (AFG3022B, Tektronix, USA). We manually the triggered stimulation when the mice were at the far-end of the open-field as to the nest position, within in a body-length distance from the wall.

### Optogenetic manipulation

Animals were handled and habituated for 10-15 min to the looming box one day before testing. During the Looming test session, mice were first allowed to freely explore the looming box for 3-5 min, then received the optogenetic manipulation or looming stimulus.

For optogenetic NpHR inhibition plus looming experiments, mice received bilateral 593 nm yellow light laser (Aurora-220-589, NEWDOON, Hangzhou) with 5-8 mW (for soma stimulation) or 15-20 mW (for terminals stimulation) light power at the fiber tips. For optogenetic ChR2 excitation plus looming experiments, mice received bilateral 10 Hz, 473 nm blue light laser (Aurora-220-473, NEWDOON, Hangzhou) with 8 mW light power at the fiber tips. Light stimulation was delivered 1 s before onset of the looming stimulus and continued until the looming was turned off. Mice received two trials of looming stimulus plus photoinhibition or photoactivation.

For optogenetic activation experiments, mice were placed into the same looming box and received a 2.5 s 473-nm blue laser (50 pulses, 20 Hz, 5 ms pulse duration) with 15-20 mW (terminal) or 5-8 mW (soma) light power at the fiber tips. No looming was presented for these experiments. Light stimulation was delivered to the SC and VTA somas, as well as the SC-VTA terminals. For the activation of VTA^GABA+^ neurons experiments, the GAD2-Cre mice received 2.5 s or 20 s blue light (60 Hz, 5 ms pulse duration, 5-8 mW) stimulation in the VTA; For the activation of VTA^DA+^ neurons experiments, the DAT-Cre mice received 2.5 s blue light (10 Hz, 5 ms pulse duration, 5-8 mW) stimulation in the VTA; no other experimental details were changed. Light was presented two times at about 3 min intervals via a manual trigger.

All light stimulation was manually presented by the experimenter when the mice were at the far-end of the open-field as to the nest position, within in a body-length distance from the wall.

For all our lost-of function experiments (optical inhibition by NpHR), the inhibition was all bilateral. For all our gain-of function experiments (optical activation of ChR2), the activation was all unilateral.

### Pharmacological antagonism

C57BL/6J mice were used for pharmacological experiments. For VTA experiment, 120 nl glutamate antagonists (0.1μg AP5, 0.001μg NBQX (2, 3-dihydroxy-6-nitro-7-sulfamoyl-benzoquinoxaline-2, 3-dione) in saline) were bilaterally injected into VTA (AP, −3.20 mm; ML, ±0.35; mm, DV: −4.5 mm). For the CeA experiment, 150nl GABA_A_ antagonist (0.005μg Bicuculline) was bilaterally injected into the CeA (AP, −1.5 mm; ML, ±2.95 mm; DV, −4.6 mm). Mice were given 2 weeks to recover after surgery. Saline (control) and antagonists were infused into the targets 30 min before a looming test to assess the antagonistic effect of the antagonism receptors.

### Behavioral analysis

Behavioral data were analyzed with Anymaze software (Stoelting Co.). Speed data was first extracted using Anymaze software and then analyzed using Matlab. Individual time courses were represented setting T-0ms as the time of stimulation. The following measures were obtained as indices of looming-evoked or light-evoked defensive behaviors: (1) latency to return nest: the time from looming stimulus or photostimulation presentation to time when the mouse escaped/entered the nest; (2) time spent in nest (% of 1 min bin): time spent in the nest following looming stimulus or photostimulation; (3) speed. The mice were allowed to move freely in the open field with a nest paradigm before looming stimulus or light stimulation. The mice were moving freely when looming stimulus or light stimulation began. Here, “baseline” was defined as the period 50 s before onset of the looming or light stimulation. The average speed during the baseline period was set as 100%. We have presented all speeds in relative percentage form compared with baseline average speed. For the speed bar graphs: post-stimulation speed was averaged over a 0.5 s-long time window centered around the time of maximum speed, detected from the time of stimulation to 10 s after.

For blinding purposes, all mice used for behavioral experiments were given a unique ear tag numerical identifier. Data obtained from mice with imprecise cannula or fiber placements were not used for analyses.

### In vivo electrophysiology

The optrode was an optic fiber (0.37 NA, 200 μm) attached around eight stereotrodes, which consisted of insulated nichrome wires (OD = 17 μm; CFW, California, USA). The tips of the stereotrode were electro-plated with platinum with platinum (chloroplatinic acid solution) until the impedance reached approximately 0.5 MΩ, and was 0.3 mm longer than the tips of fiber. Data were collected by OmniPlex D Neural Data Acquisition System (Plexon, Dallas, USA). Mice were anesthetized with urethane (10% W/V, 1.9 g/kg body weight, i.p.) and positioned in the stereotaxic apparatus. The optrode was placed into the VTA (AP, −3.20 mm; ML, −0.3 mm; DV, – 4.1~4.7 mm). If the detected signal in the VTA was stable, a 473 nm laser (20 Hz with 5 ms width pulses) was delivered to excite CaMKIIα-positive projections from the SC. Baseline was recorded in the VTA for 2-3 min, and a 2.5 s optic stimulation was delivered at 1 min intervals. At the end of the experiment, electrolytic lesions were performed (0.1 mA DC, 20 s) to label the recording side.

### Data analysis

Recorded spikes were detected and sorted by using Plexon Offline Sorter software (Plexon, Inc., Dallas, TX, USA), then analyzed in Neuroexplorer (Nex Technologies, Madison, AL, USA) and Matlab (MathWorks, Natick, MA, USA). Putative dopamine neurons were classified by agglomerative clusters from linkages based on the following electrophysiological criteria (Wang and Tsien, 2011a): 1) low baseline firing rate (<10 Hz); 2) inter-spike interval (ISI)>4 ms within a baseline firing rate (<10 Hz); 2) inter-spike inter(AP) widths broader ⩾ 1) ms. In contrast, if neurons did not meet these criteria, they were classified as non-putative DA neurons, VTA neurons were recorded for around 3 min to establish the discharge properties and basal firing rate of VTA neurons before optical stimulation. The baseline firing rate was calculated using the mean and standard deviation (SD) of firing rate values for every 0.5 s bin 2.5 s preceding optical stimulation. To determine the excitatory and inhibitory response related to photostimulation, we used peristimulus time histograms (PSTHs) to analyze firing pattern (Wang and Tsien, 2011a). 1) Excitation: the PSTH 20 ms (0.5 ms per bin) following the light pulses was calculated. The neuronal responses were defined as the 3 consecutive bins after the onset of light stimulation triggered the PSTH that exceeded the 2 SDs of baseline value. 2) Inhibition: the PSTH in 10 s (0.2 s per bin) following the light pulses was calculated. Neuronal inhibition was defined as the 5 consecutive bins after onset of light stimulation triggered the PSTH that dropped to at least 35% below baseline value. The magnitude of the normalized firing rate was calculated according to the following equation: Magnitude = (counts in response period)-(mean counts per bin in baseline) x (number of bins during the response period).

### Fiber Photometry

The fiber photometry system (ThinkerTech, Nanjing) consisted of a 480 nm excitation light from a LEDs (CREE XPE), reflected off a dichroic mirror with a 435–488 nm reflection band and a 502–730 nm transmission band (Edmund, Inc.), and coupled into a 200 μm 0.37 NA optical fiber (Thorlabs, Inc.) by an objective lense. The laser intensity at the fiber tip was about 20 μW. GCaMP6s fluorescence was collected using the same objective, transmitted by the dichroic mirror filtered through a green fluorescence protein (GFP) bandpass emission filter (Thorlabs, Inc. Filter 525/39), and detected by the sensor of an CMOS camera (Thorlabs, Inc. DCC3240M). A Labview program was developed to control the CMOS camera which recorded calcium signals at 50 Hz. The behavioral event signal was recorded by a DAQ card (NI, usb-6001) at 1000 Hz using the same program.

### Data analysis

All the raw data were smoothed with a moving average filter (5 point span) and then segmented and aligned according to the onset of looming stimulus within individual trials or bouts. The fluorescence change (ΔF/F) values were calculated as (F-F_0_)/F_0_, where F_0_ is the baseline fluorescence signals averaged over a 2 s-long control time window (typically set 1 s) prior to a trigger event.

To compare activity between different conditions, bar graphs were computed by averaging data along a 0.5 s time window centered around the time of the activity pic. Time courses were made by averaging individual trials aligned to the time of stimulation. A multivariate permutation (1000 permutations, α level of 0.05) test was used to account for data significance level on time courses, and a threshold indicating statistically-significant increase from the baseline was applied (p<0.005). Areas surrounding the time courses and error bars represents mean ± SEM.

## QUANTIFICATION AND STATISTICAL ANALYSIS

The number of biological replicates in each group was 3-6 mice per group for anatomy, 4-8 mice for in vitro and in vivo physiology, 5–7 mice per group for fiber photometry, and 5-16 mice per group for behavior. These numbers were based on previously published study (Wei et al., 2015; Tovote et al., 2016). Data distribution was assumed to be normal, but this was not formally tested. All statistics were performed in Graph Pad Prism (GraphPad Software, Inc.), unless otherwise indicated. Paired student test, unpaired student test, one-way ANOVA and two-way ANOVA were used where appropriate. Bonferroni post hoc comparisons was conducted to detect significant main effects or interactions. In all statistical measures a P value <0.05 was considered statistically significant. Post hoc significance values were set as *P< 0.05, **P< 0.01, ***P< 0.001 and ****p< 0.0001; all statistical tests used are indicated in the figure legends.

## ACKNOWLEDGMENTS

We thank Minmin Luo for providing us with GAD2-ires-Cre mice, Zilong Qiu for TH-Cre mice and Yangling Mu for DAT-ires-Cre mice. We thank Cornelius T Gross for comments on our manuscript. We also thank Wei He, Tiaotiao Liu and Mi Xia for conducting the electrophysiology analyses.

This work was supported by National Natural Science Foundation of China (NSFC) 31630031 (L.W.), NSFC 81425010 (L.W.), NSFC 31471109 (L.L), NSFC 31671116 (J.T.), NSFC 91632303/H09 (F.X.), and NSFC 31500861 (P.W.); International Partnership Program of Chinese Academy of Sciences 172644KYS820170004 (L.W.); External Cooperation Program of the Chinese Academy of Sciences GJHZ1508 (L.W.); Guangdong Provincial Key Laboratory of Brain Connectome and Behavior 2017B030301017 (L.W.); Shenzhen governmental grants JCYJ20150529143500959 (L.W.), KQJSCX20160301144002 (L.L), JSGG20160429184327274 (L.L), JCYJ20150401150223647 (Z.Z.), JCYJ20160429190927063 (J.T.) and JSGG20160429190521240 (F.Y.); Shenzhen Discipline Construction Project for Neurobiology DRCSM [2016]1379 (L.W.); Ten Thousand Talent Program (L.W.). Guangdong Special Support Program (L.W.).

## CONTRIBUTIONS

Z.Z. and X.L. contributed equally to this work. Z.Z., X.L, and L.W. designed the project and Z.Z. and X.L initiated the project, performed virus and/or drug injections, photometry recordings, optogenetic behavior testing, electrophysiology experiments, and collected and analyzed the data. S.C. contributed to whole cell patch clamp recording experiments. Z. Zhang and X.H preformed rabies virus injections and immunohistochemistry. M.Q. contributed to analysis of the photometry data. Y.L., Y.T. and C.C. helped collect the data. P.W. and N.L contributed to pilot experiments. L.L., G.B., C.G., G.F., F.X., and L.W interpreted the results and commented on the manuscript. Z.Z., X.L., L.L., and L.W. wrote the manuscript. L.W. and F.X. supervised all aspects of the project.

